# Identification of non-coding elements associated with the evolutionary loss of digit 1

**DOI:** 10.64898/2025.12.17.695027

**Authors:** Jacob A. Scott, Anastasiia Lozovska, Jessica Perez, Martin J. Cohn

## Abstract

Evolutionary loss of digits and limbs occurred many times during the diversification of tetrapods. The limbs of the earliest tetrapods were polydactylous, bearing as many 8 digits, but after the pentadactyly (five digits) was stabilized, no lineage evolved more than five digits per limb but many underwent digit loss. Within mammals, digit 1 (thumb on the forelimb or hallux on the hindlimb) is lost most frequently and is typically the first digit to disappear. Evidence from numerous studies of limb development shows that digit 1 development is controlled by mechanisms that are at least partially independent of the posterior 4 digits. In addition, evolutionary loss of digit 1 occurred independently in forelimbs and hindlimbs. Despite the frequency of digit loss in mammals, the underlying genetic mechanisms have not been uncovered. We used comparative genomics and bioinformatics to reconstruct the evolutionary history of genomic changes associated with digit 1 loss in the forelimbs and/or hindlimbs of 65 mammalian species. We identified a set of conserved non-coding elements that evolved in concert with independent (parallel) losses of digit 1. These elements, which are frequently near genes with known roles in digit development, underwent accelerated evolution in digit-reduced lineages. Furthermore, we find that elements with accelerated rates of evolution show significant loss of binding sites for transcription factors that are crucial for anterior digit development. The results suggest that *cis*-regulatory elements important for the development of digit 1 are evolutionarily accelerated in mammalian lineages that underwent reduction and loss of digit 1.

## Introduction

Extant tetrapod species have a maximum of five digits on each limb (known as pentadactyly), but many species have fewer than five digits due to evolutionary losses. Indeed, digit loss has occurred independently in at least one lineage of every major class of tetrapods. Reduction and loss of digits appears to evolve independently between the forelimb and the hindlimb and generally occurs gradually, one digit at a time (McHorse et al. 2019; Smith et al. 2020)(Solounias et al. 2018). Which digits are lost first in a given taxa is related to developmental timing. The “last in-first out” principle describes the phenomenon in which the last digit to be specified during development is, in general, the first digit to be lost in evolution. For example, salamanders and mammals differ in the order of digit development (the last digit to form is digit 5 in the former and digit 1 in the latter) and the first digits to be lost in these groups tend to be the last to develop, irrespective of its identity (Royle and Young 2021; Trofka et al. 2021) (Fröbisch and Shubin 2011). Digit 1 has been lost or vestigialized (which will be collectively referred to as “digit reduction” hereafter) independently at least 17 times in the mammalian forelimb and at least 14 times in the mammalian hindlimb (Senter and Moch 2015).

Reduction of digit 1 in mammals can occur without affecting the remaining four digits, suggesting that patterning, chondrogenesis, and development of digit 1 can be modulated independent of digits 2 through 5. In the forelimbs of anthropoid primates, for example, variation in the length of the first digit is independent of the posterior digits, which scale with the distal radius (Reno et al. 2008). Consistent with this evolutionary modularity, the molecular mechanisms of digit development differ between digit 1 and the posterior digits. For example, digits 2-5 show combinatorial expression of *Hoxd10-13*, whereas only a single Hoxd gene, *Hoxd13*, is expressed in digit 1 (Montavon et al. 2008; Reno et al. 2008). Moreover, digit 1 is the only digit that develops independent of *Shh* (Chiang et al. 2001; Ros et al. 2003). Studies of digit development across a range of tetrapods suggest that digit 1 may be the only digit with a conserved molecular identity through all amniotes (Stewart et al. 2019).

Despite the prevalence of digit 1 loss on either forelimbs or hindlimbs during mammalian evolution and developmental evidence of a partially independent digit 1 module, isolated reduction of digit 1 is rarely observed in human or mouse mutants. For example, in human congenital anomalies, loss of digit 1 is typically associated with a broader syndrome (Van De Laar et al. 2007; Jain et al. 2011; Vanlerberghe et al. 2019; Nguyen and Ho 2022). Similarly, in the mouse *Hoxa13^Hd^* mutant, reduction of digit 1 in forelimbs and hindlimbs is accompanied by anomalies in digit 5 and the carpals/tarsals (Fromental-Ramaint et al. 1995). Indeed, a mutant is yet to be discovered that perfectly recapitulates the digit 1 reduction phenotypes that are observed in nature.

Most developmental control genes studied to date are involved in development of multiple anatomical features, a concept known as pleiotropy (reviewed in Stearns 2010). Given that mutations in pleiotropic genes can have widespread effects, evolution has favored modifications to regulatory elements that control tissue-specific activity of pleiotropic genes over changes to coding sequences themselves (reviewed in Carroll 2008). Thus, if the evolutionary losses of digit 1 resulted from mutations in non-coding elements that regulate gene expression in the anterior region of the autopod, then this might explain why gene knockouts have not recapitulated the isolated loss of digit 1.

The regulatory mechanisms that underlie evolutionary modularity of digit 1 are not well understood, and, as a result, a number of questions remain unresolved. If loss of digit 1 occurred by regulatory evolution, then do the causal elements govern known digit patterning networks or do they act on genes whose functions in digit development are yet to be determined? Have the many independent losses of digit 1 during the diversification of mammals evolved by modulation of the same pathways, or did the same phenotype evolve by different mechanisms in different lineages? Moreover, does forelimb- or hindlimb-specific loss of digit 1 in mammals reflect independent genetic control of pollux (thumb) and hallux (great toe) development?

Here, we address these questions using a comparative genomics approach to study 65 species that show partial or complete loss of digit 1 in the forelimbs and/or hindlimbs. Analysis of the relationships between digit 1 reduction and the substitution rates of conserved non-coding elements (CNEs) throughout the genome revealed patterns of accelerated evolution in genomic elements with potential roles in digit 1 development in forelimbs and hindlimbs. Accelerated elements were found more frequently than expected near genes associated with digit reduction pathologies in humans. In many cases, these elements are forelimb- or hindlimb-specific, which could explain the independent reduction of digit 1 between the limbs. Finally, we show that these elements evolve frequently in highly conserved regions and involve disruptions to similar transcription factor binding sites (TFBS), suggesting that accelerated evolution of CNEs could have direct effects on transcription of genes that regulate development of digit 1 in a forelimb- and hindlimb-specific manner.

## RESULTS

### Digit 1-reduction is evident prior to interdigital apoptosis

To identify genomic elements associated with evolutionary losses of digit 1, we examined the genomes of 94 mammalian species that have either retained or reduced digit 1 on forelimbs and/or hindlimbs. The survey spanned 15 orders and included 47 species that retain digit 1 in both the forelimbs and the hindlimbs, 4 species that underwent reduction of the thumb (pollux; digit 1 on the forelimb), 14 species that underwent reduction of the hallux (digit 1 on the hindlimb), and 29 species with reduction of both the thumb and hallux (Fig. 1 and supplementary table S1). In identifying these phenotypes, we also sought embryonic limb phenotypes from digit-reduced species using publicly available and published datasets (Canine Embryonic Atlas, Baker Institute for Animal Health, Cornell University, https://www2.vet.cornell.edu/canine-atlas; Dettlaff and Vassetzky 1991; Acker et al. 2001; Knospe 2002; Cunha et al. 2005; Franciolli 2011; Cooper et al. 2014; Lopez-Rios et al. 2014; Tissières et al. 2020; Butler and Juurlink). In each case that we examined, reduction of digit 1 was apparent at the early digit condensation stage, before the onset of interdigital apoptosis (Fig. 1c).

**Fig. 1.**
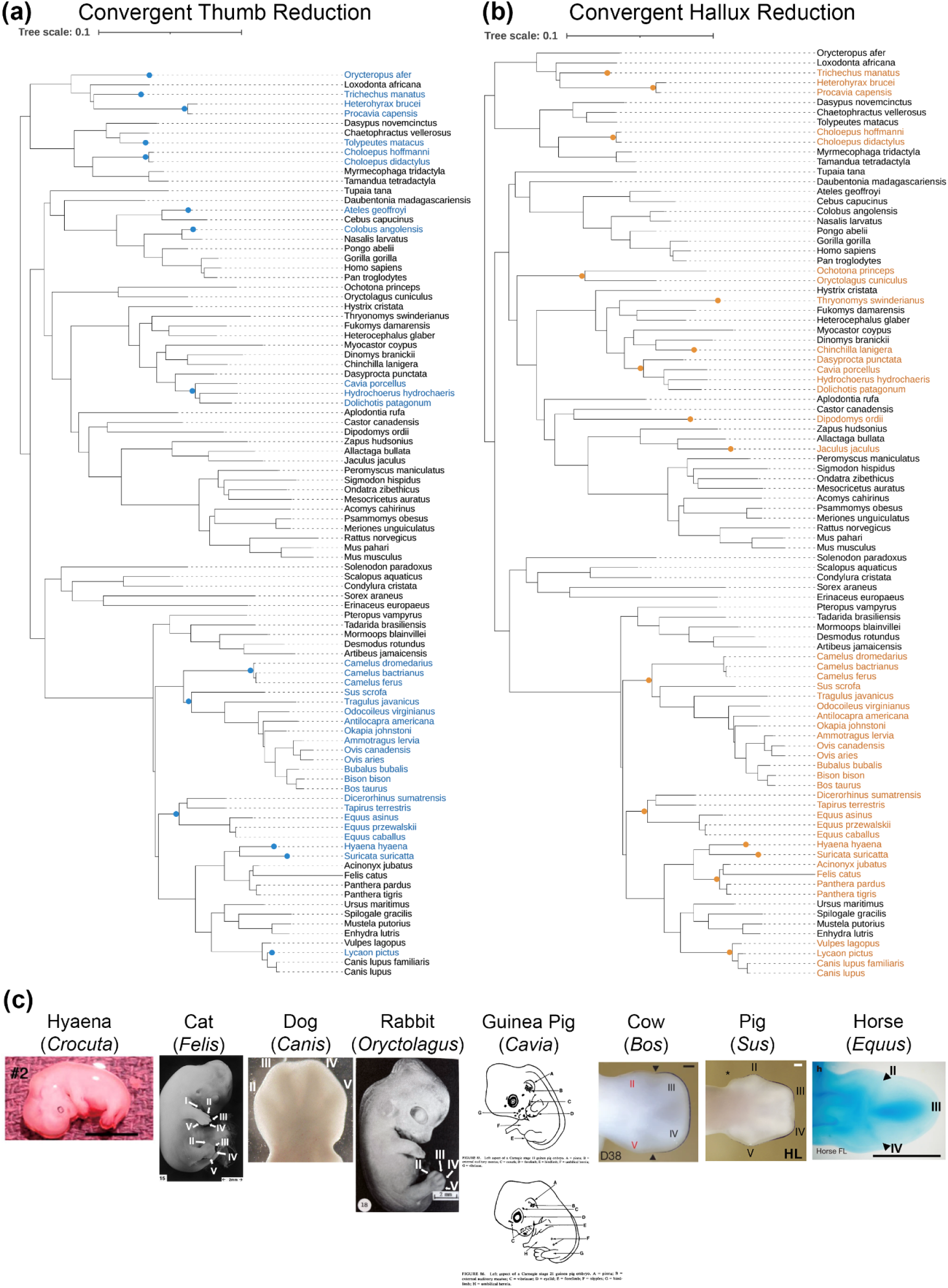
Cases of convergent digit 1 reduction used in this study. a) Species with thumb reduction highlighted in blue; origin of thumb reduction event indicated by blue circle. Species in black text retain their hallux, regardless of the phenotype of their other digits. b) Species with hallux reduction highlighted in orange; origin of hallux reduction event indicated by orange circle. Species in black text retain their hallux, regardless of the phenotype of their other digits.

### Digit 1-reduced species show accelerated evolution of elements near genes involved in human digit anomalies

Next, we searched for genomic regions that are conserved among species that retain digit 1. We identified 441,637 genomic elements that are conserved in species that retain their thumbs and 431,188 genomic elements that are conserved in species that retain their halluces. We then tested these elements for signatures of accelerated evolution in species that underwent evolutionary reduction of digit 1 on the forelimbs or hindlimbs.

#### Thumb loss elements

Species that exhibit reduction of digit 1 in the forelimb show accelerated evolution of 3,486 conserved elements. Of these, 2,171 elements overlapped ENCODE’s predicted cis-regulatory elements (CREs) and/or ChromHMM predicted limb enhancers (referred to hereafter as “thumb-loss CREs”). Analysis of the 2,171 thumb-loss CREs using the Genomic Regions Enrichment of Annotations Tool (GREAT) showed overrepresentation of these elements around genes associated with human congenital diseases that involve reduction of digits (Fig. 2a), validating the ability of this strategy to recover functionally relevant candidates. This subset of accelerated elements was associated with 2,375 genes (Fig. 2c). The majority of these elements were located either 0-5kb or 50-500kb from the transcription start site of their associated gene (Fig. 2b). The three most enriched Gene Ontology (GO) Biological Processes for this dataset were “Anterior/posterior pattern specification”, “Regulation of chromatin organization”, and “Endochondral bone morphogenesis.” These data also show enrichment of the human phenotype terms “Short middle phalanx of finger” and “Aplasia/hypoplasia of the middle phalanges of the hand” (Fig. 2a).

**Fig. 2.**
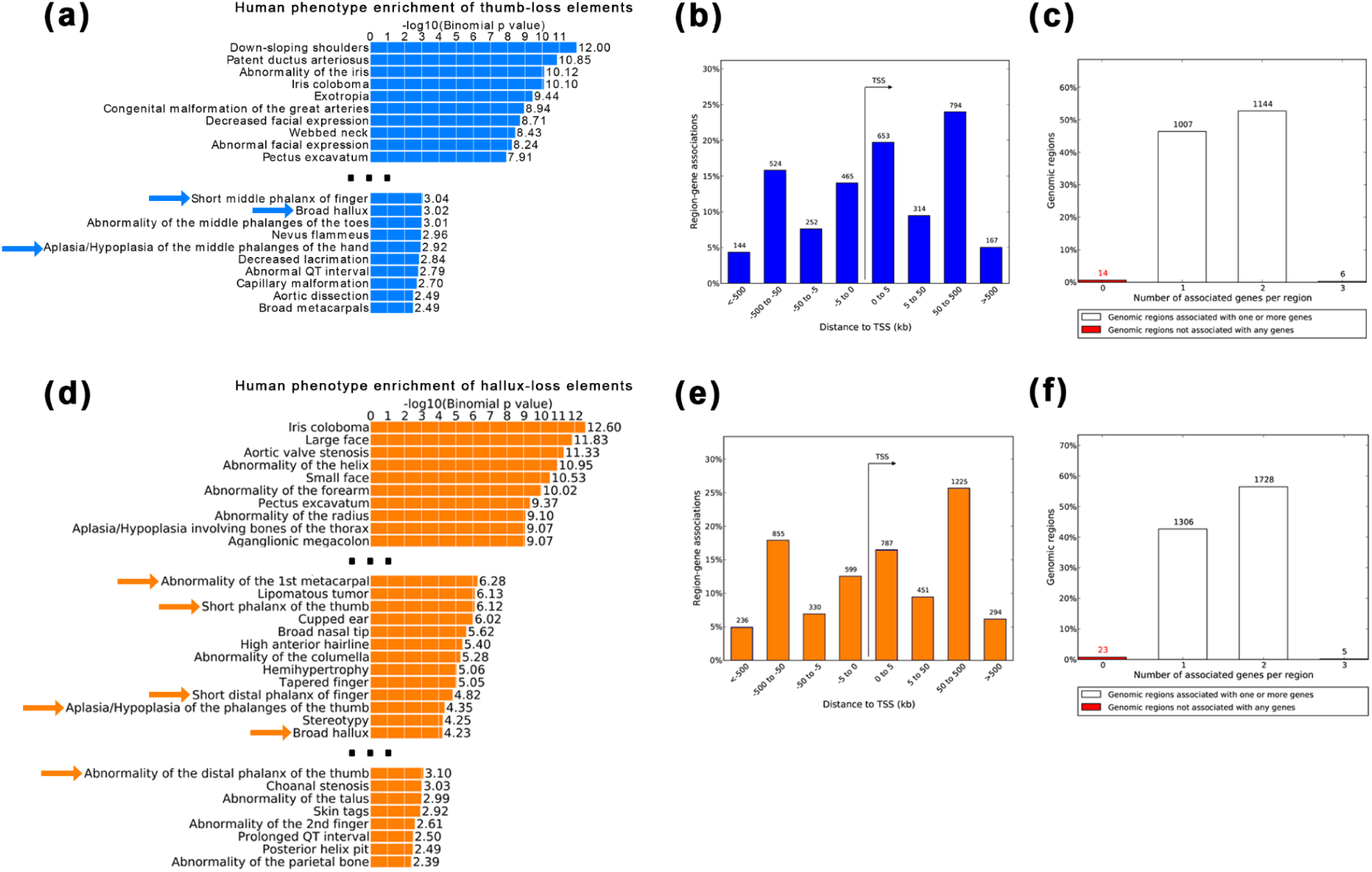
a,d) Human phenotype terms enriched by thumb- (a) and hallux- (b) loss elements. Includes the top 10 most enriched terms, with breaks (…) until digit reduction-related terms. Blue (thumb) and orange (hallux) arrows indicate terms relevant to digit reduction or abnormalities of digit 1. b,e) The distance of each thumb- (b) or hallux- (e) loss element from the transcription start sites of their associated genes. c,f) Number of genes associated with thumb- (c) and hallux- (f) loss elements.

These thumb-loss CREs overlapped at least partially with eight Vista enhancers that show activity in the E11.5 mouse limb (Visel et al. 2007; Kosicki et al. 2025)(supplementary table s2). Two of these enhancers (hs1431.0 and hs305.0) drive expression in the anterior domains of the mouse forelimb at stage E11.5. Four of these enhancers (hs1262.0, hs142.0, hs1462.0, and hs2672.0) drive expression in proximal domains of the mouse limb at stage E11.5. One enhancer (hs2951.0) drove expression at low levels throughout the entire E11.5 mouse embryo.

#### Hallux loss elements

In species that have undergone reduction of digit 1 in the hindlimb, we identified 4,885 conserved elements that show accelerated evolution. Of these, 3,062 elements overlapped ENCODE’s predicted CREs and/or ChromHMM predicted limb enhancers (referred to hereafter as “hallux-loss CREs”). GREAT analysis of the 3,062 hallux-loss CREs showed overrepresentation around genes involved in human congenital diseases that involve digit reduction, including some that specifically involve the reduction of digit 1 (Fig. 2d).

These elements were associated with 2,994 different genes (Fig. 2f). As with the thumb-loss elements, the majority of hallux-loss elements were either 0-5kb or 50-500kb from the transcription start sites of their associated genes (Fig. 2f). The two most enriched GO Biological Processes were “Anterior/posterior pattern specification” and “Negative regulation of ossification.” These data also were enriched for the human phenotype terms “Abnormality of the 1st metacarpal”, “Short phalanx of the thumb”, “Aplasia/hypoplasia of the phalanges of the thumb”, and “Abnormality of the distal phalanx of the thumb”, among other less specific digit-related terms (Fig. 2d).

Hallux-loss CREs overlap at least partially with 12 Vista enhancers shown to be active in the E11.5 mouse limb, 4 of which were also identified as thumb-loss enhancers (supplementary table s2). Two of these enhancers (hs1431.0 and hs2724.0) drive expression in the anterior of the mouse hindlimb at stage E11.5. Four of these enhancers (hs142.0, hs1603.0, hs1681.0, and hs1745.0) drive expression in the proximal domains of the E11.5 mouse hindlimb. Three of these enhancers (hs1471.0 and hs313.0) drive expression in posterior domains of the E11.5 mouse hindlimb. One of these enhancers (hs2951.0) drives expression at low levels throughout the E11.5 mouse embryo. The remaining three enhancers (hs1339.0, hs2121.0, and hs2585.0) drive expression only in the forelimb and not in the hindlimb of E11.5 mice.

### Thumb-specific and hallux-specific elements show the strongest associations with human digit 1-loss phenotypes

The majority of D1-loss elements were accelerated in either the forelimbs (thumb-loss) or hindlimbs (hallux-loss). Of the 2,171 thumb-loss and 3,062 hallux-loss elements that we identified, only 1,063 overlapped genomic coordinates between the two datasets (Fig. 3a). Outside of these shared elements were 1,108 thumb-specific elements and 2,002 hallux-specific elements.

**Fig. 3.**
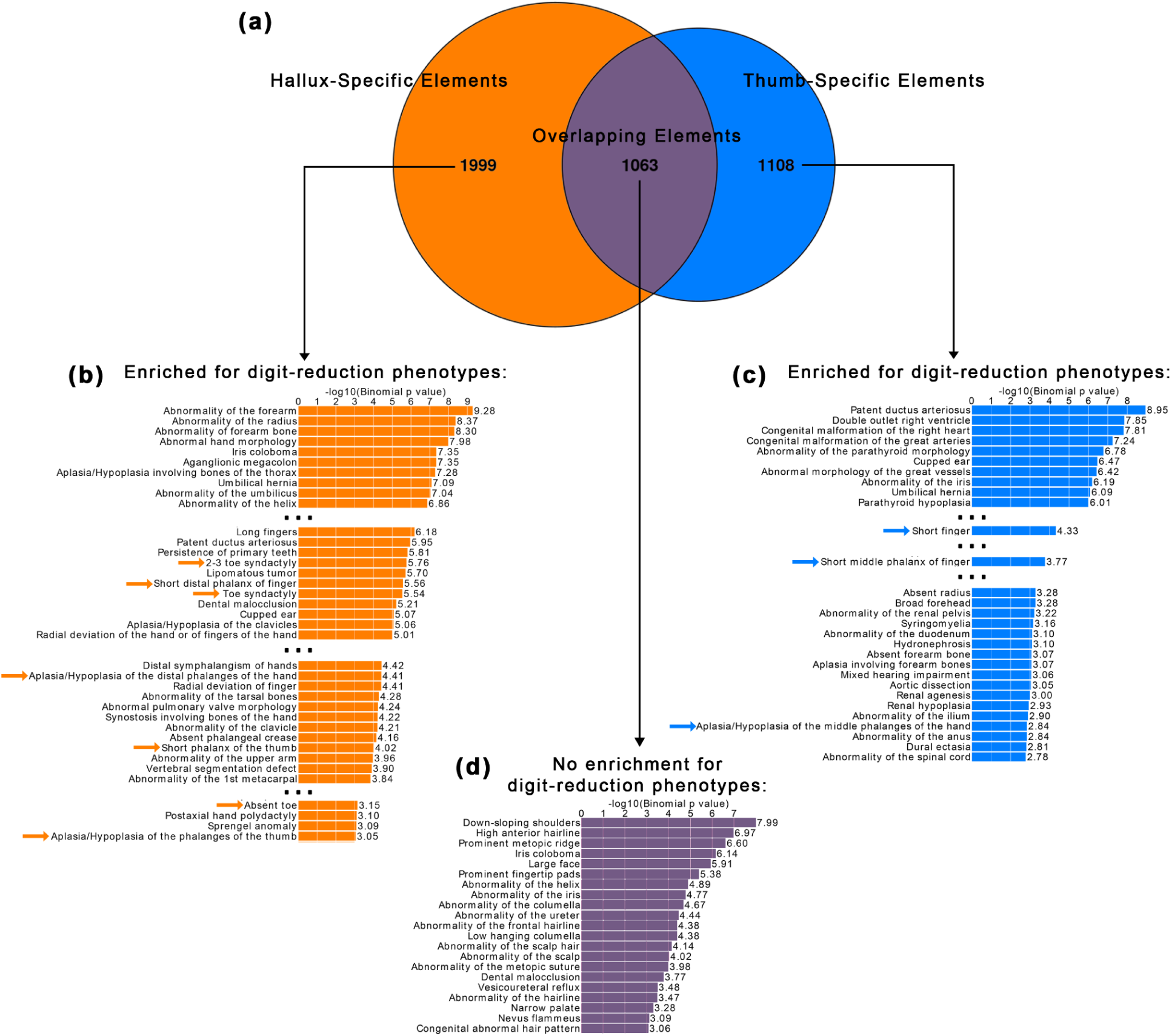
a) Number of thumb- and hallux-loss elements that do and do not overlap genomic coordinates. b-c) Human phenotype terms enriched by thumb- (b) and hallux- (c) specific elements. Includes the top 10 most highly enriched terms, as well as any terms that pertain to abnormalities of the digits or anterior limb. d) Human phenotype terms enriched by elements with overlapping coordinates between thumb- and hallux-loss elements. No digit reduction phenotypes were observed in this analysis.

When thumb-specific, hallux-specific, and overlapping elements were re-analyzed for enrichment of ontological terms, both the thumb-specific and hallux-specific elements were enriched for digit reduction terms (Fig. 3b,c). For example, the thumb-specific elements were enriched for “Aplasia/hypoplasia of the middle phalanges of the hand” and the hallux-specific elements were enriched for “Aplasia/hypoplasia of the phalanges of the thumb.” By contrast, overlapping elements (i.e. those shared between the thumb and hallux datasets) showed no significant enrichment for any digit reduction phenotypes (Fig. 3d). The only limb-related term enriched in the overlapping elements was “Prominent fingertip pads.” This suggests that the majority of CREs involved in digit 1 development/loss may be forelimb- or hindlimb-specific.

### Convergent acceleration of elements around genes involved in anterior digit development

We next tested for hotspots of accelerated evolution by examining mammalian lineages that underwent parallel evolution of digit 1 loss. We defined digit 1 loss “hotspots” as genes associated with thumb- or hallux-specific elements in at least one third of the lineages with digit 1 loss (considering thumb- and hallux-loss hotspots separately).

We identified 183 thumb-loss hotspots and 440 hallux-loss hotspots, of which 43 were present in both datasets (**table 1**). Some genes had nearby acceleration in the majority of digit 1 loss lineages, but no gene had nearby acceleration in every lineage, suggesting that, despite some degree of convergence, there are multiple paths to evolve digit 1 loss. Additionally, different lineages often showed acceleration of different enhancers around the same gene, suggesting that there are multiple ways to modulate expression of the same digit 1 patterning genes.

**Table 1.**
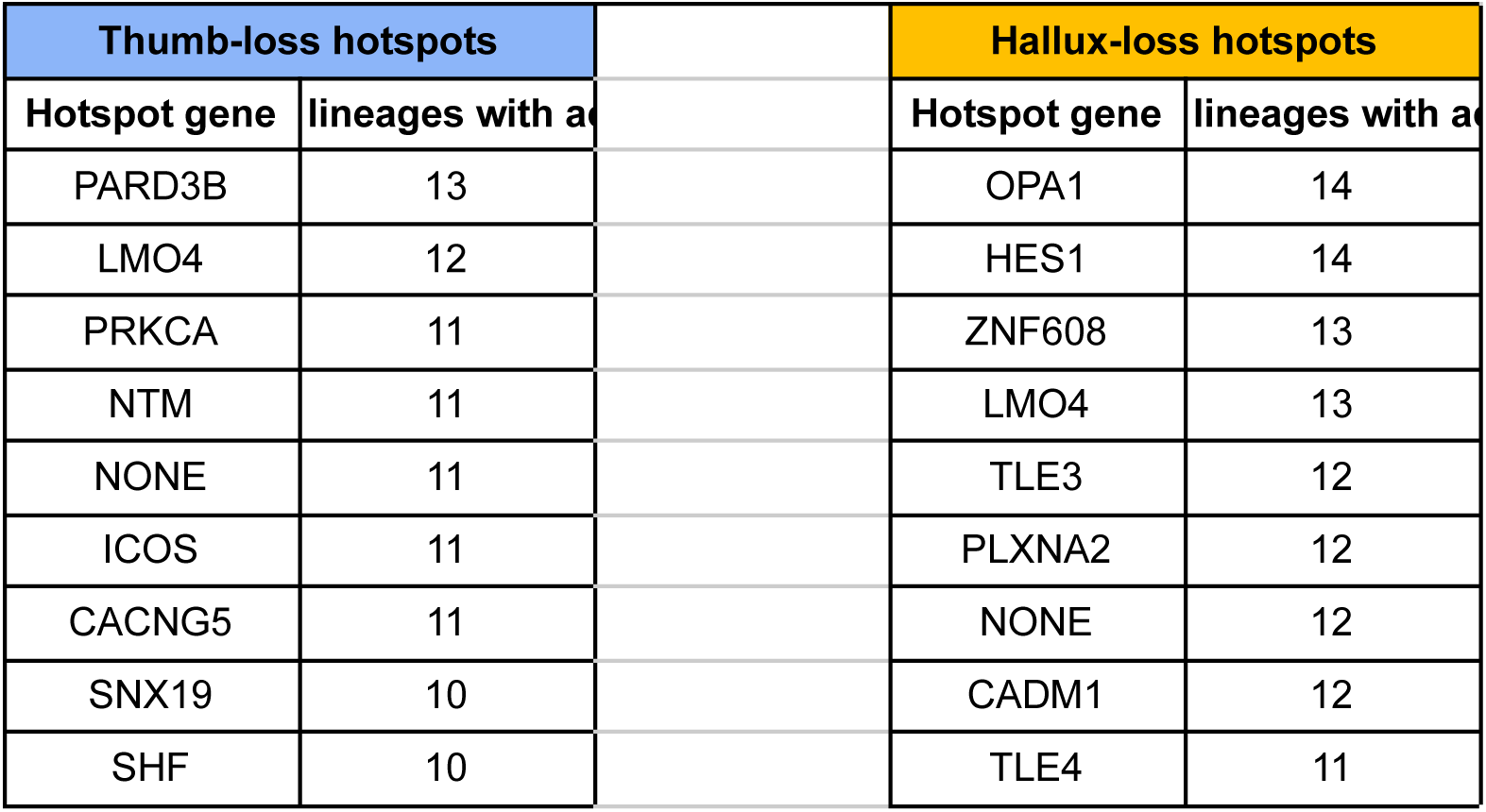
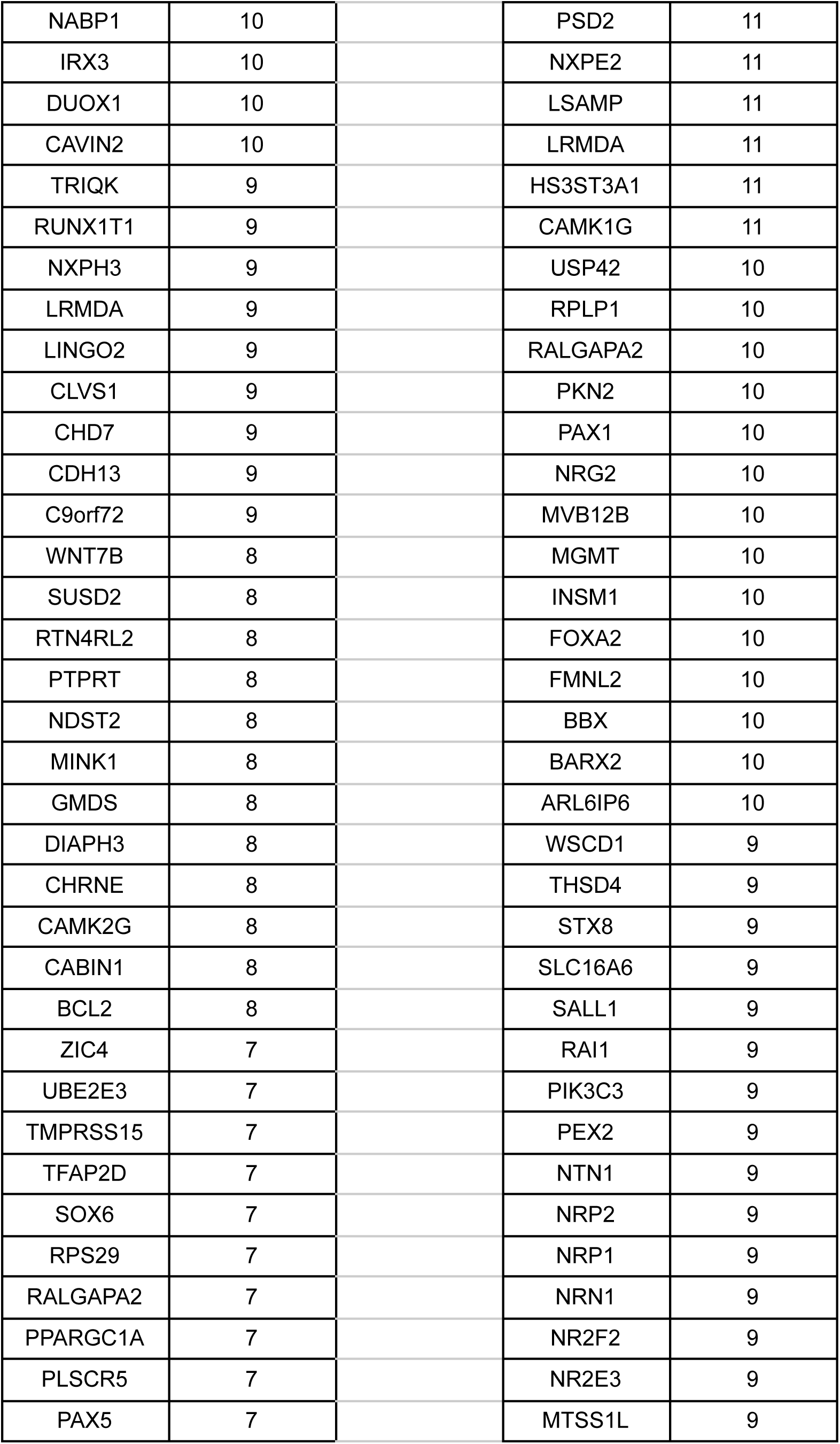
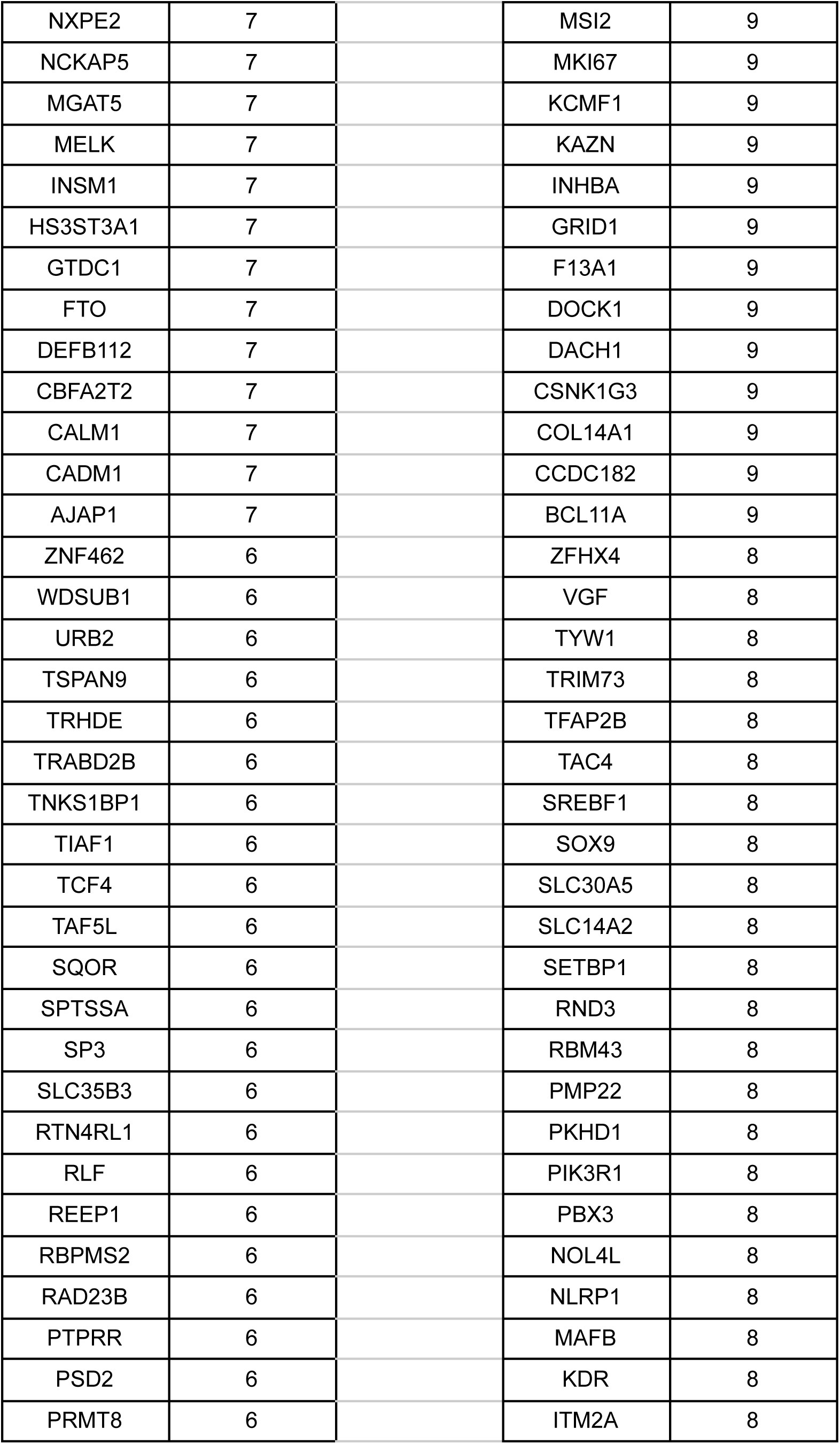
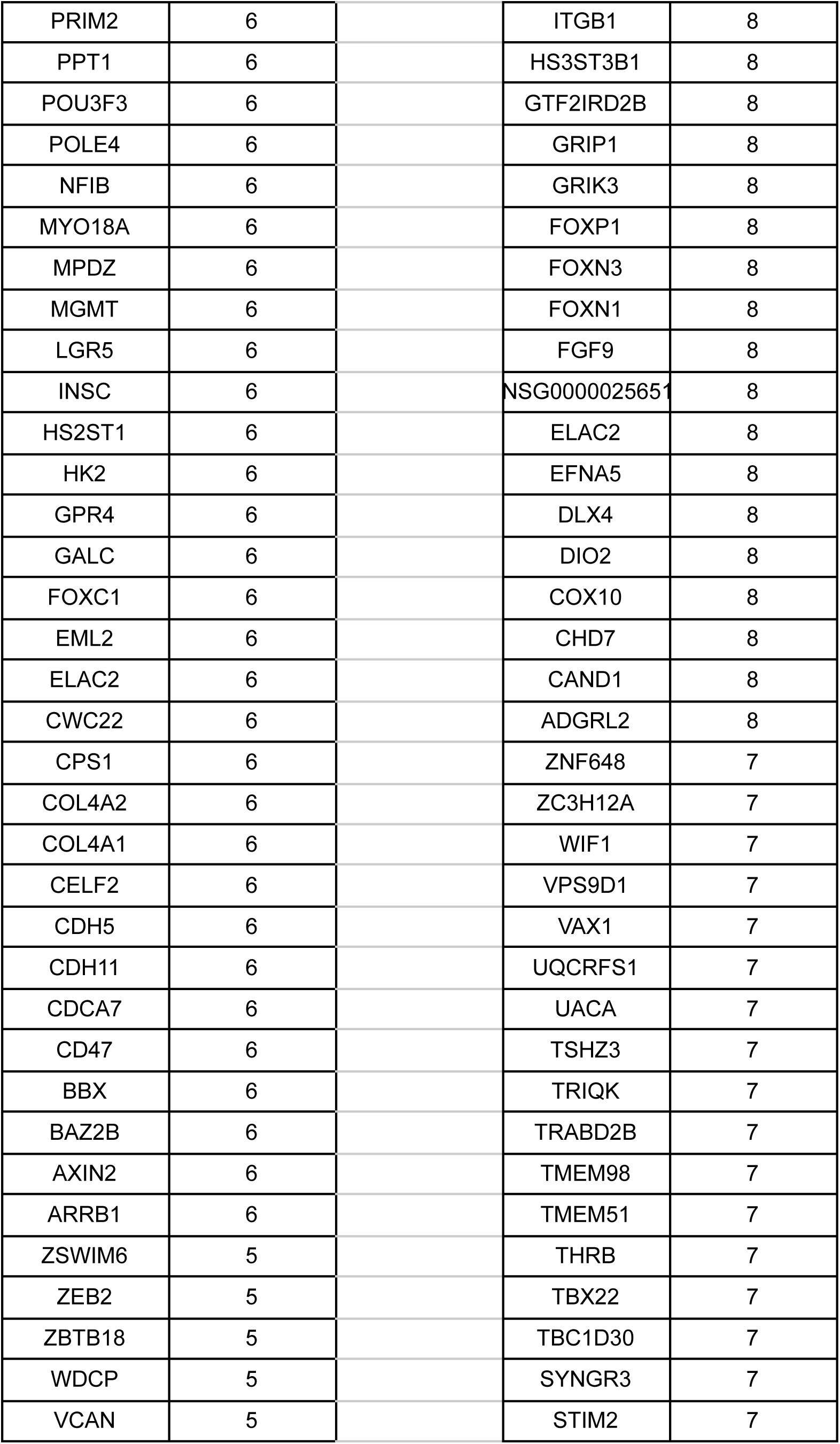
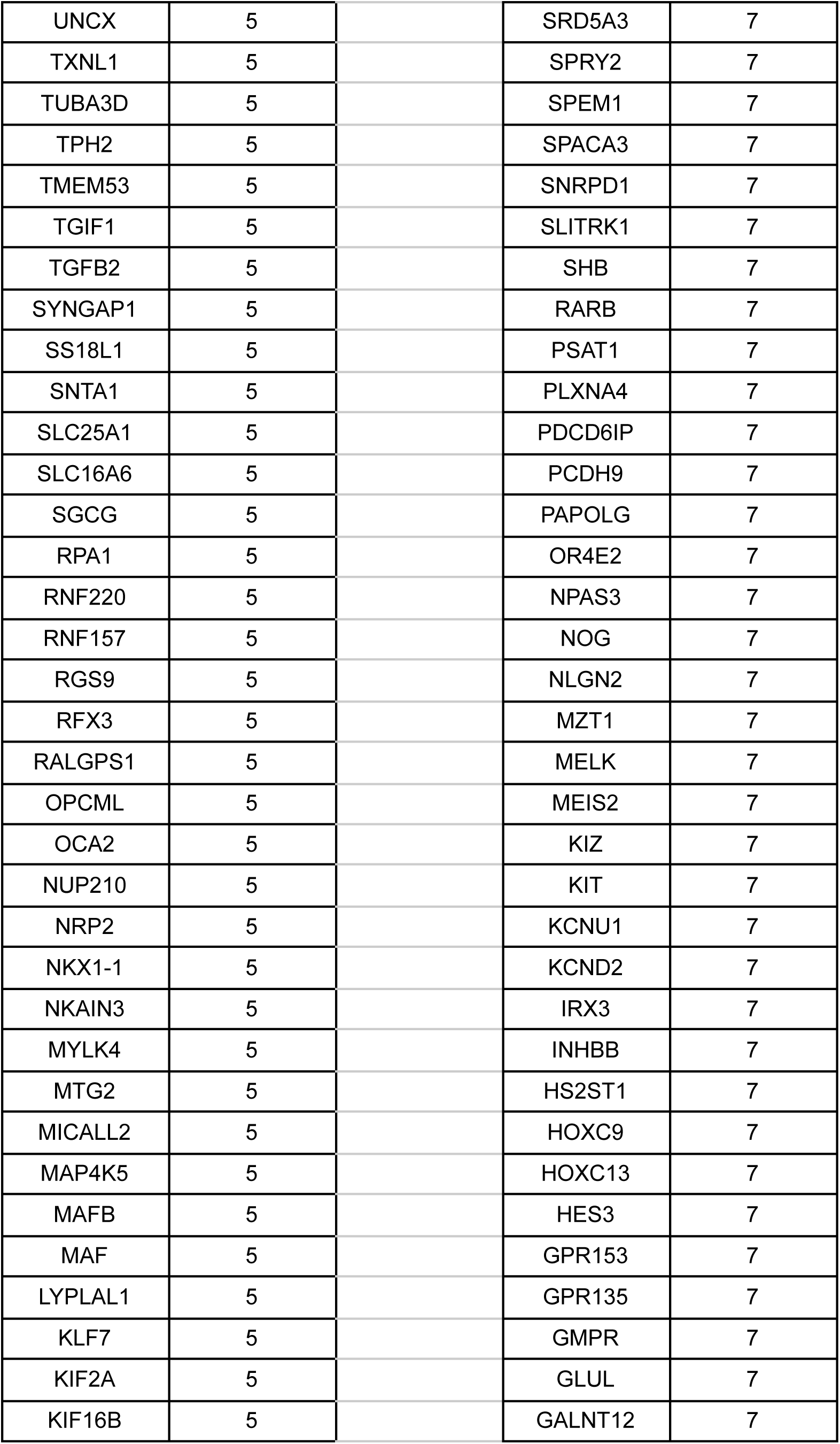
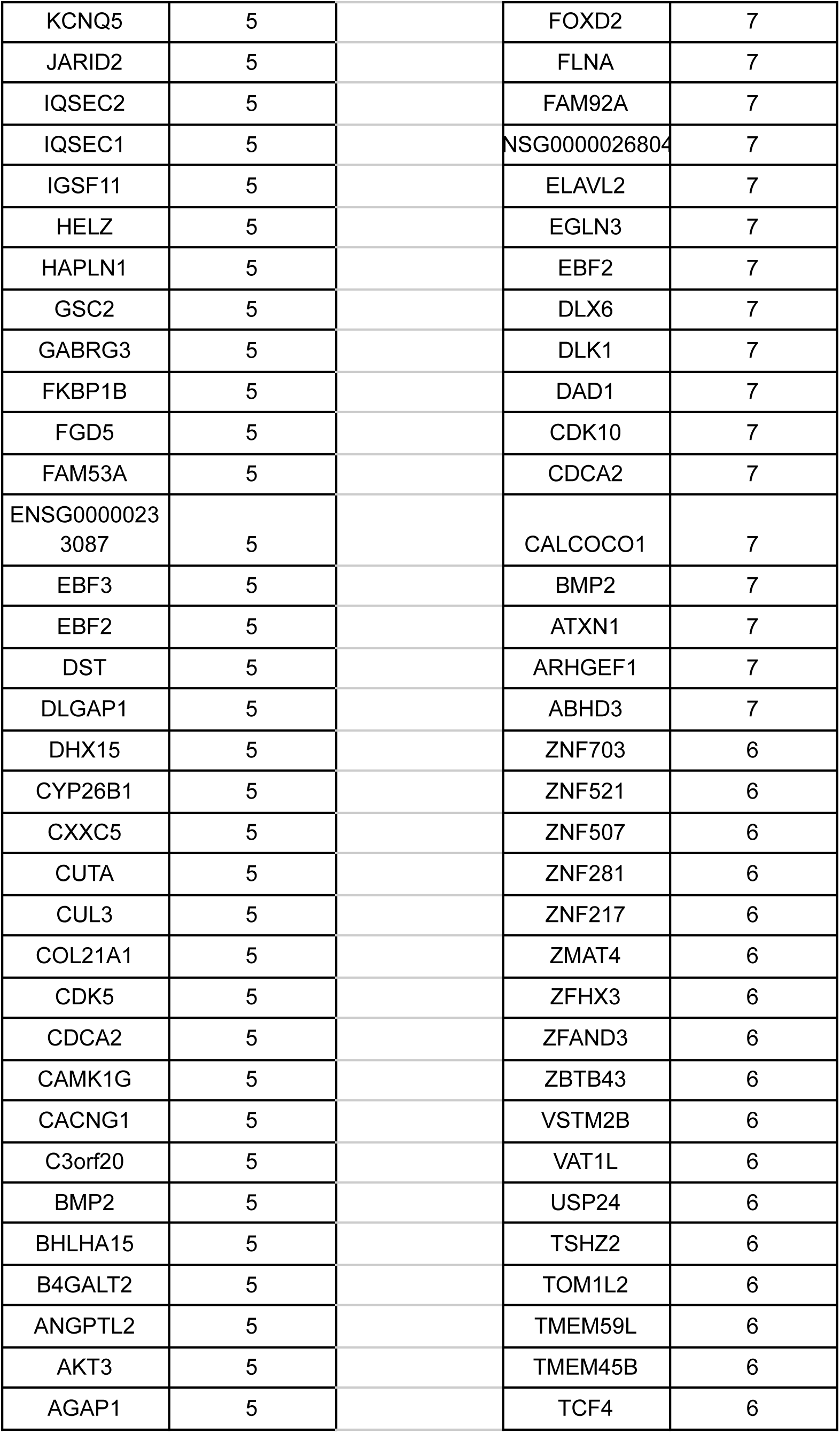
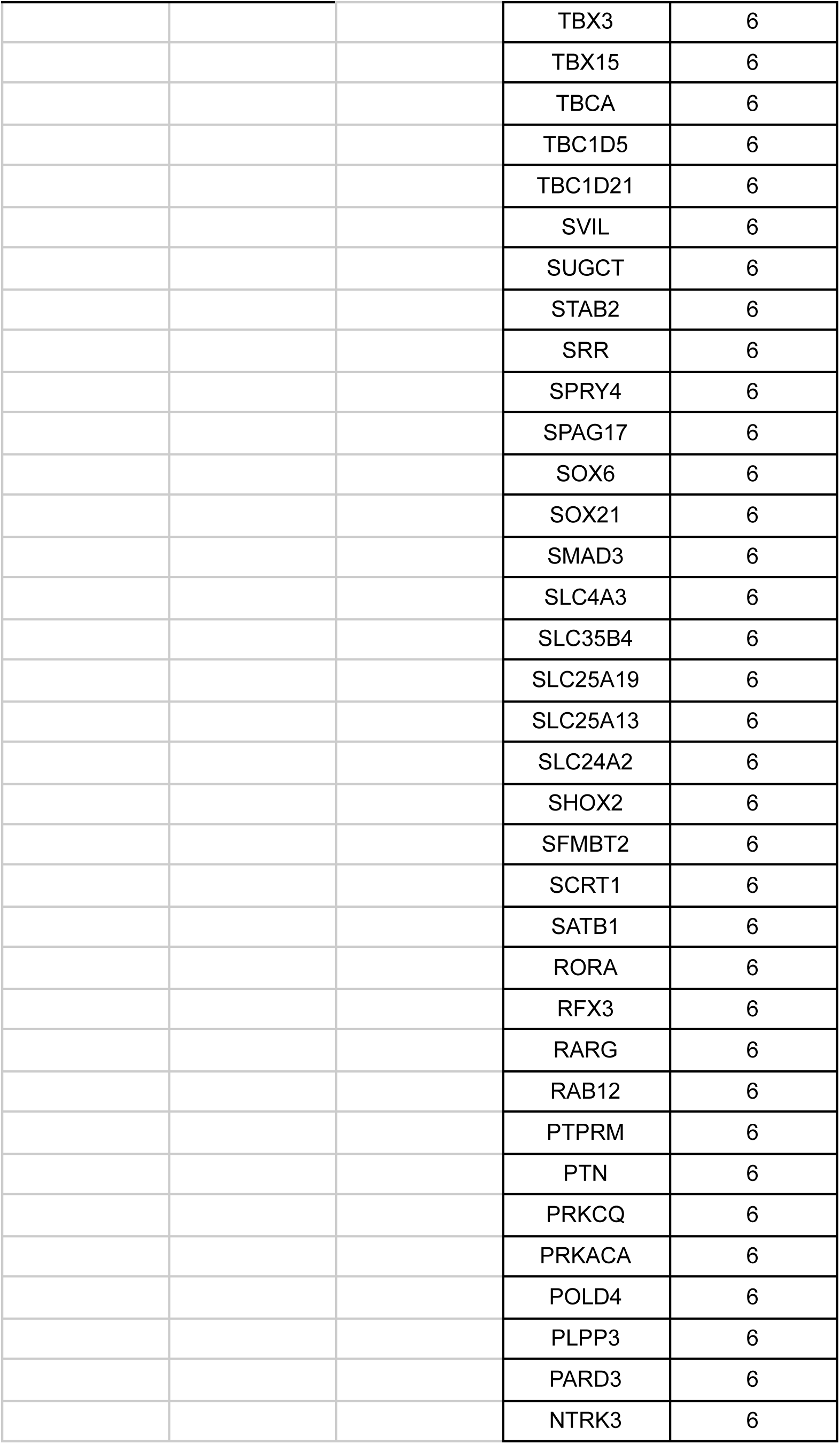
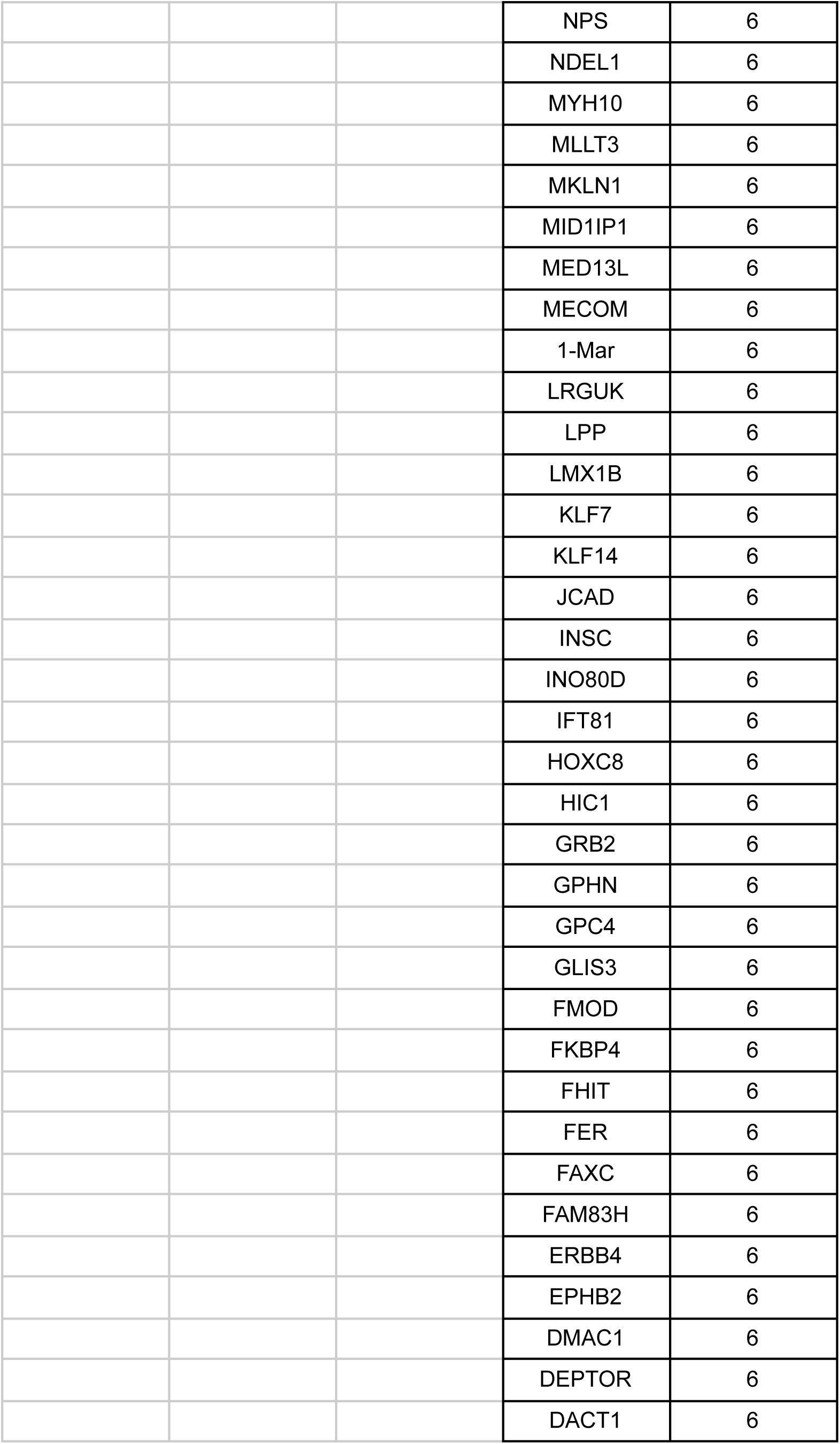
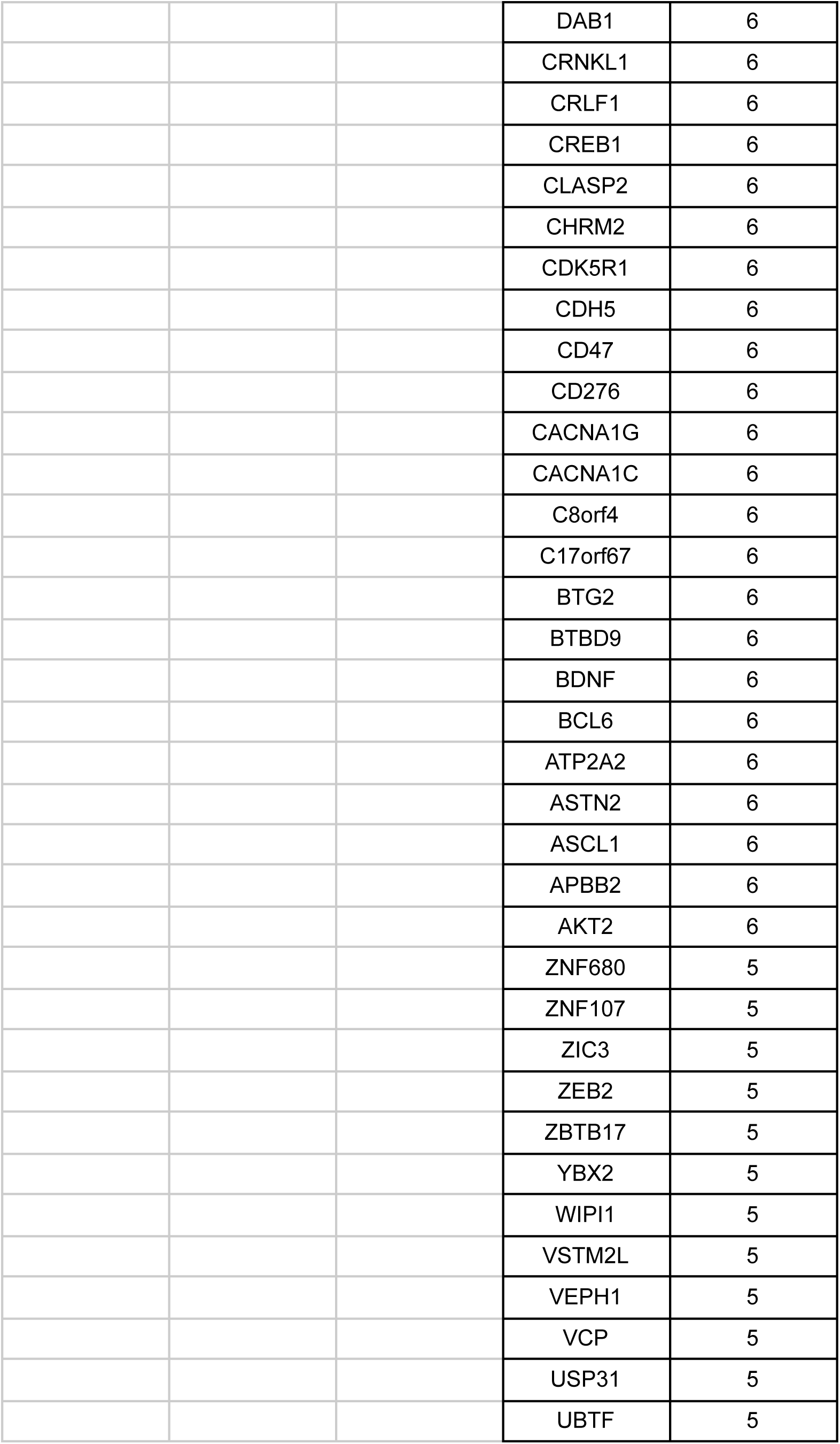
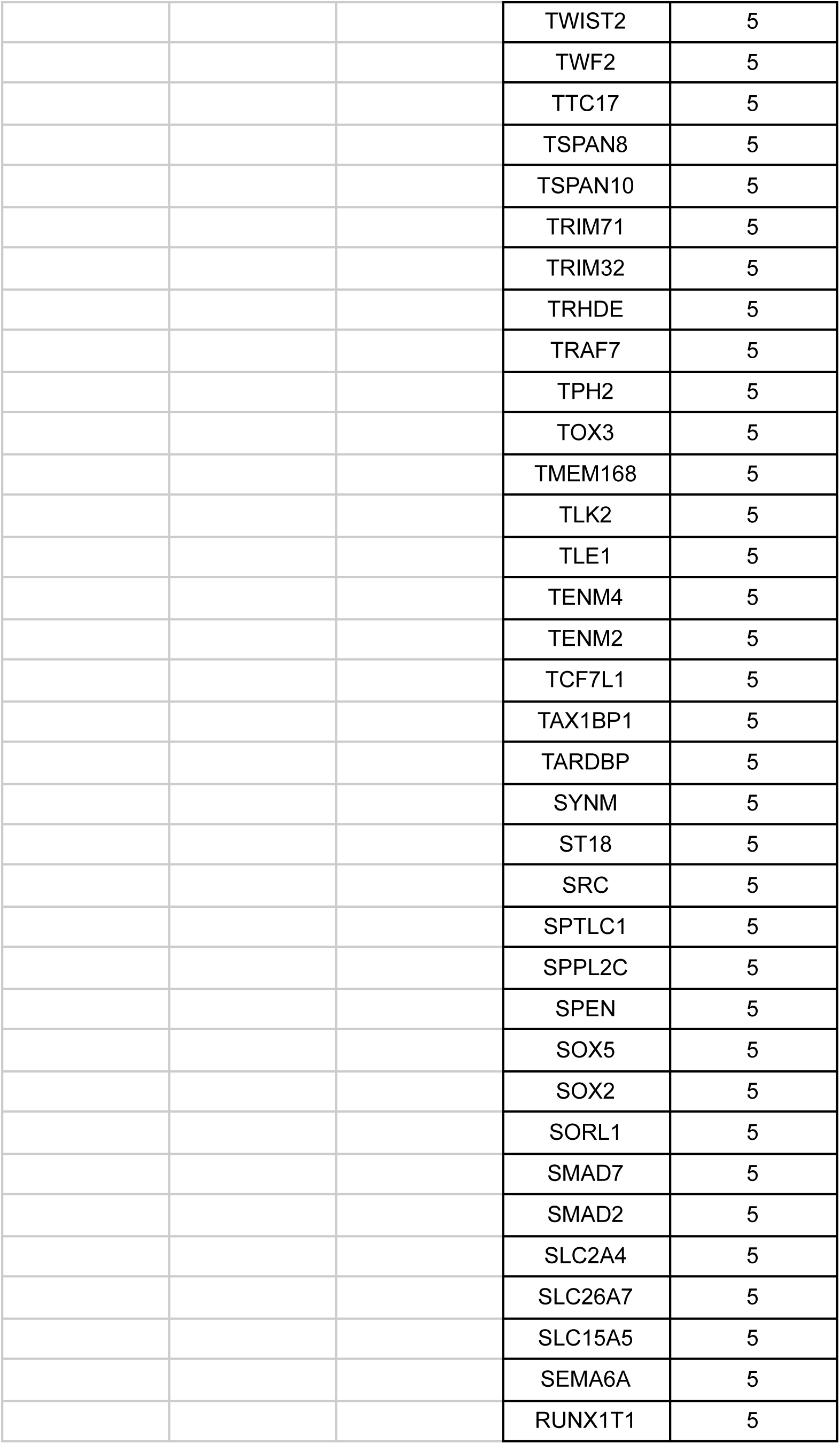
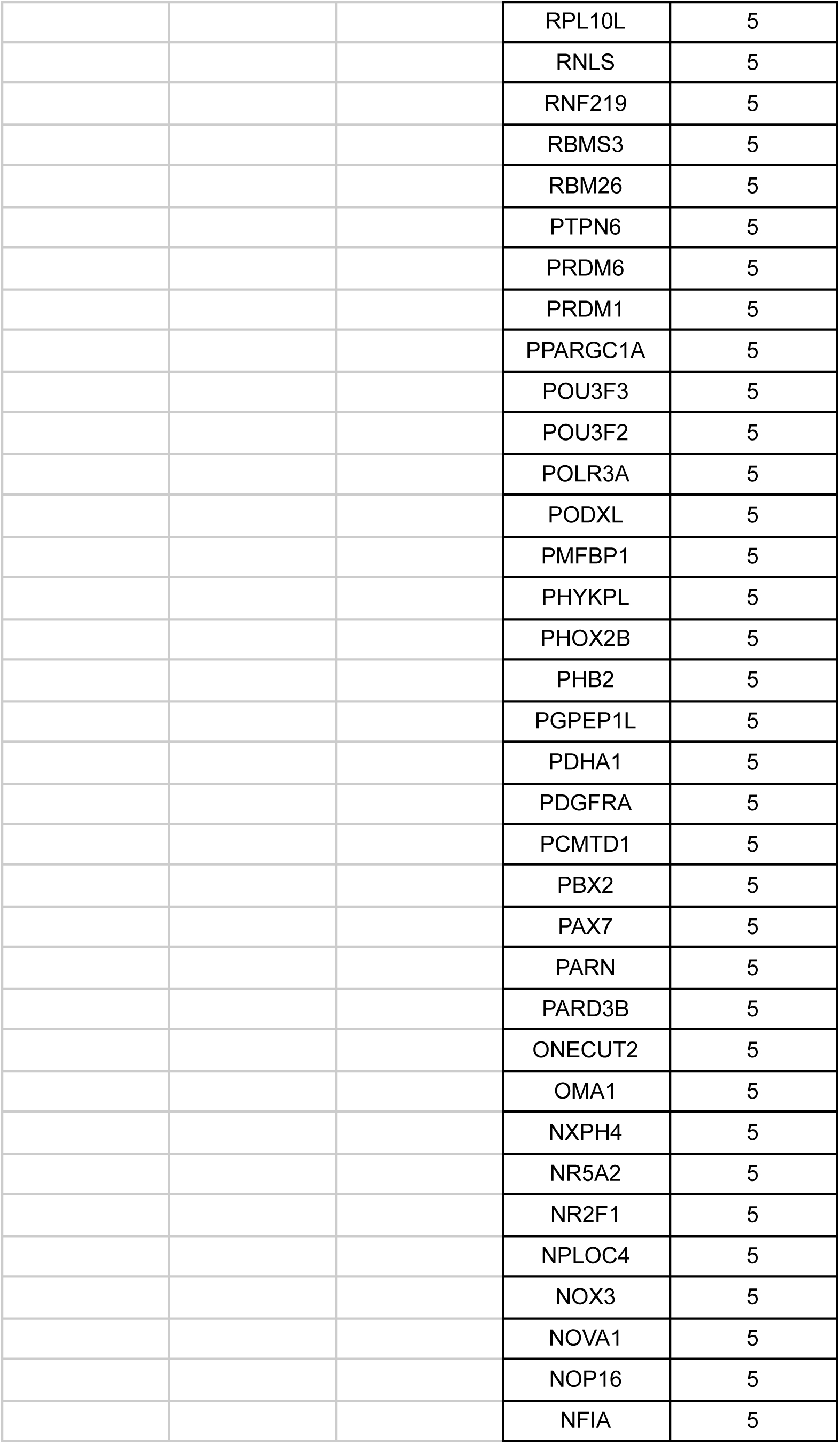
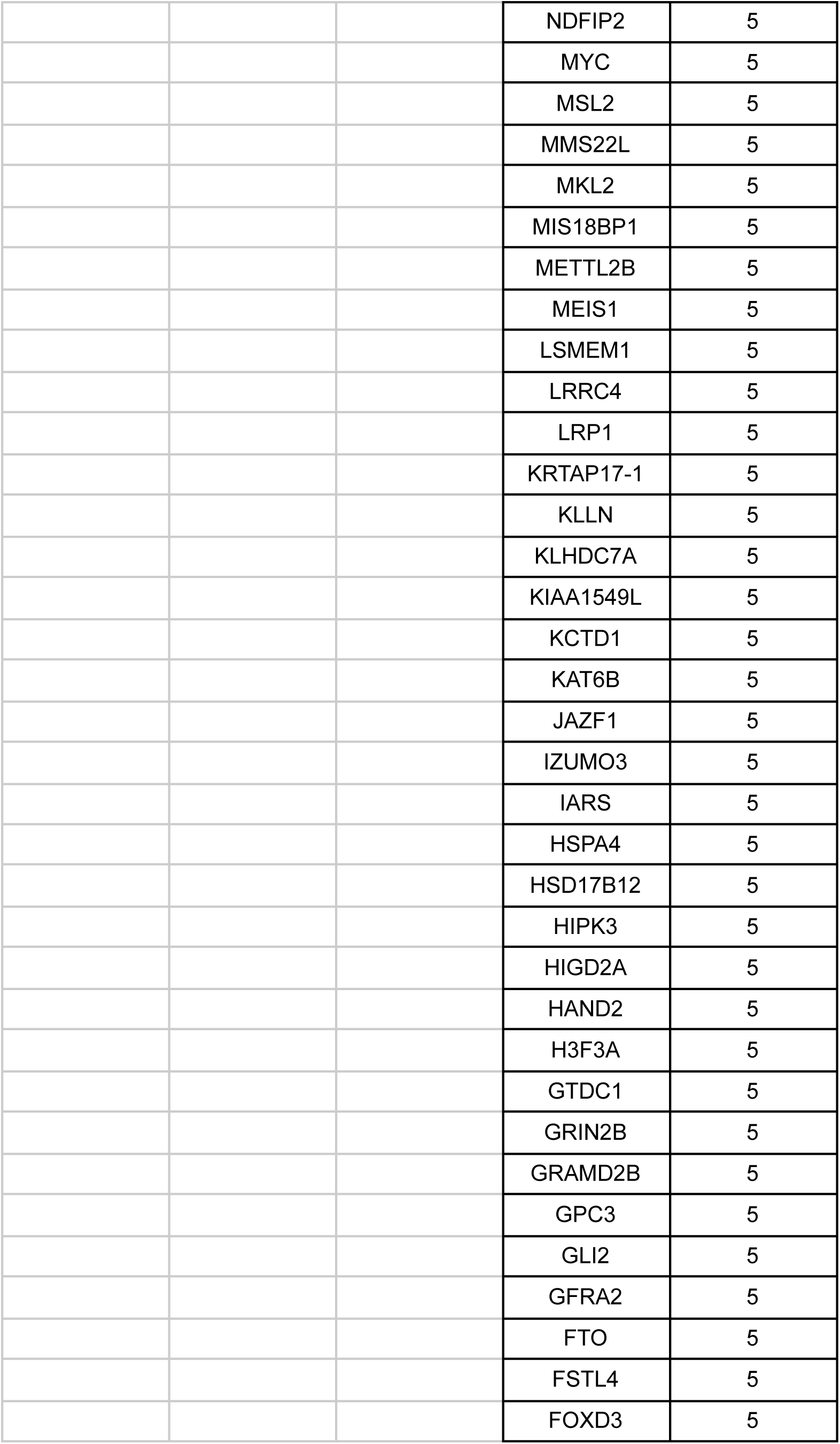
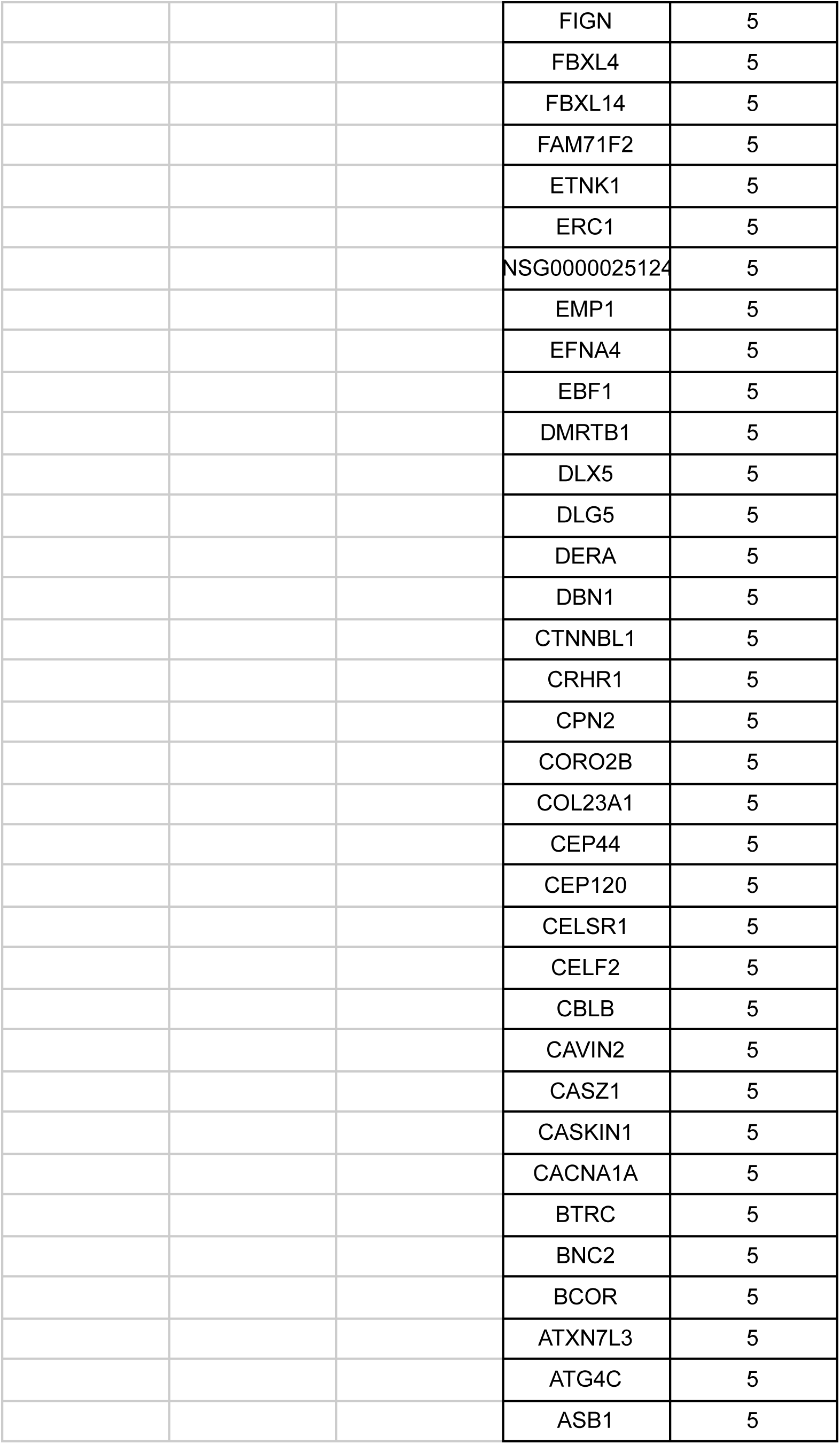
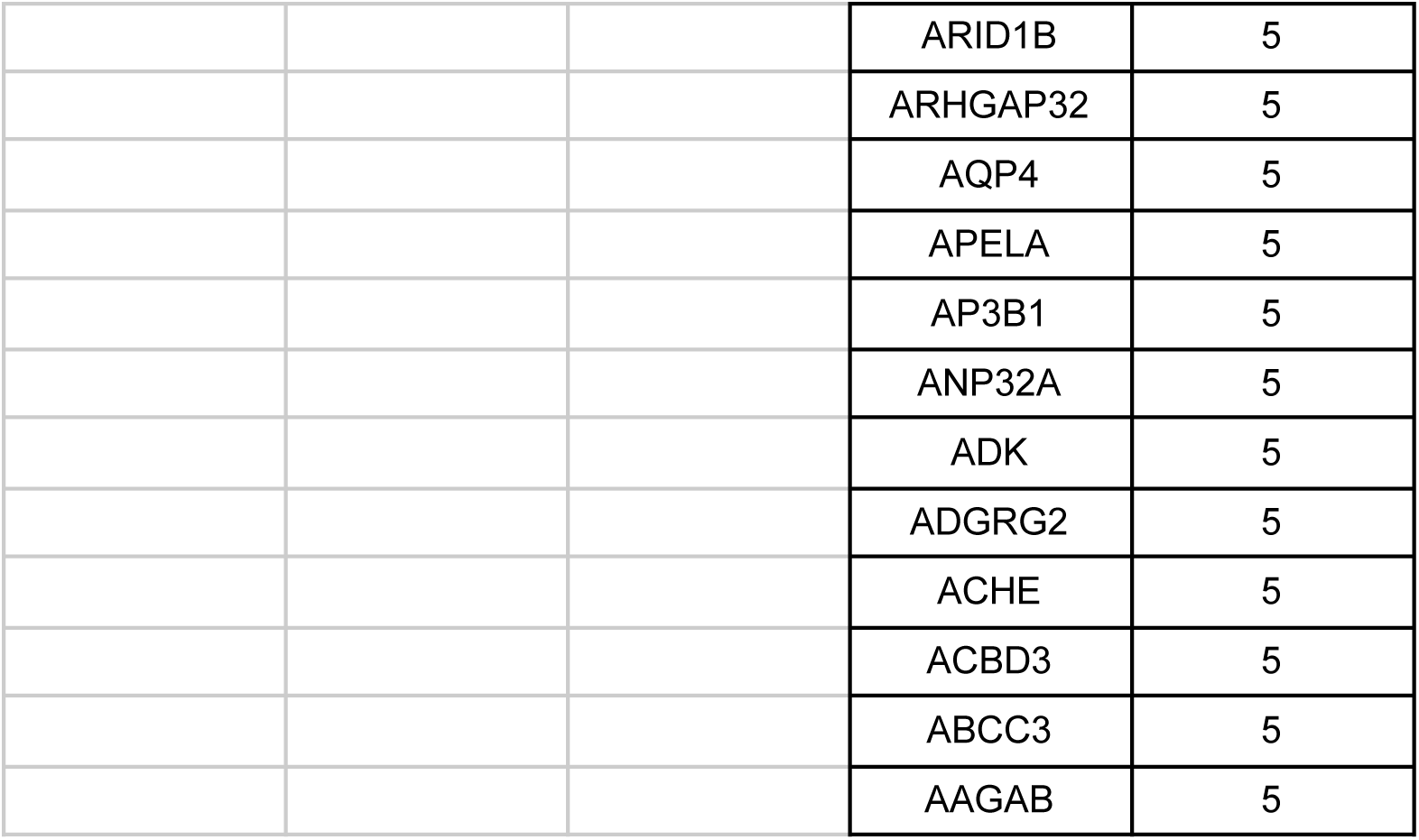
List of thumb- and hallux-loss hotspots.

Among the most frequently accelerated thumb-loss hotspots is the region around *Irx3*, where acceleration occurred in 10 of the 14 lineages analyzed. *Irx3* is known to be an anterior patterning gene in limb development (Yokoyama et al. 2017), and double knockouts of *Irx3* and *Irx5*, which are separated by only 600kb in the mouse IrxB cluster, show loss of anterior skeletal limb elements, including the hallux and the tibia (Li et al. 2014). The 4 taxa that do not show acceleration of elements near *Irx3* are *Lycaon* (a carnivore), *Ateles* (a primate), Caviidae (rodents), and *Trichechus* (a siren), indicating that *Irx3*-independent thumb-loss is not a phylogenetic phenomenon.

*Irx3* is also a hallux-loss hotspot, showing accelerated elements in 7 out of 15 lineages. Relative to thumb loss, hallux loss evolves around a more diverse suite of anterior patterning genes, including *Sall1* (accelerated in 9 out of 15 lineages), *Pbx3* (8 out of 15 lineages), *Dlx6* (7 out of 15 lineages), *Zic3* (5 out of 15 lineages), and *Dlx5* (5 out of 15 lineages)(Di Giacomo et al. 2006; Kawakami et al. 2009; Vieux-Rochas et al. 2013; Li et al. 2025). Fourteen of 15 lineages – artiodactyls being the exception – show acceleration near at least one of these anterior patterning genes. Hotspots also were detected near *Tfap2b* (8 out of 15 lineages) and *Hand2* (5 out of 15 lineages), which have been shown to be differentially expressed between the developing digit 1 and the rest of the autopod across amniotes (Stewart et al. 2019). Among the most frequent hotspots of hallux loss are genes in the Notch signalling pathway, including *Hes1* (14 out of 15 lineages), *Tle3* (12 out of 15 lineages), and *Tle4* (11 out of 15 lineages). *Hes1* overexpression results in polydactyly of digit 1 (Sharma et al. 2021), but the roles of *Tle3* and *Tle4* in limb development are unknown at present.

Notably absent from either list of hotspots are *Hoxa13*, *Hoxd13*, and *Gli3*, which play critical roles in development of digit 1 (Bastida et al. 2020). Indeed, only 2 lineages have accelerated elements near any of these genes; one hallux-loss *Gli3* element is accelerated in *Dipodomys* (kangaroo rat) and *Procaviidae* (hyrax), and a second hallux-loss *Gli3* element is accelerated only in *Dipodomys*. Taken together, these results indicate that parallel loss of digit 1 in different mammalian lineages is associated with evolutionary acceleration at common loci.

### Identification of anteriorly biased expression of *Lmo4*

The region surrounding LIM domain gene *Lmo4* is a strong hotspot for both thumb- and hallux-loss (accelerated in 12 out of 14 thumb-loss lineages and 13 out of 15 hallux-loss lineages). Although little is known about the function of *Lmo4* in limb development, *Lmo4* has previously been reported to be expressed in the dorsal mesenchyme of the E11.5 mouse limb (Kenny et al. 1998), though the sections presented in that study do not provide a global view of the limb. Therefore, we investigated the expression of *Lmo4* in forelimbs and hindlimbs of mouse embryos at E11.5 and E12.5, the early stages of digit development.

In E11.5 forelimbs, *Lmo4* is expressed in a U-shaped domain of the autopod that spans the perimeter of the mesenchyme, with the strongest expression detected in the anterior region of the handplate (Fig. 4a). In hindlimbs at the same stage, *Lmo4* expression was detected in 2 distinct domains of the autopod, one in the anterior mesenchyme and one in the distal mesenchyme (Fig. 4b). At E12.5, *Lmo4* is expressed throughout digit 1 and along the anterior edge of digit 2, with faint signal detected in the regions surrounding the other digits. This pattern is stronger in the forelimb (Fig. 4c) but also can be seen in the hindlimb (Fig. 4d). Thus, *Lmo4* expression is enriched in the anterior region of the autopod and becomes increasingly restricted to the region where digit 1 develops as chondrogenesis progresses. Comparison of these data with *Lmo4* expression in human limbs (Human Embryonic Limb Atlas (Zhang et al. 2024)) shows that the anterior bias of *Lmo4* is conserved between rodents and primates with pentadactyl digit formulae.

**Fig. 4.**
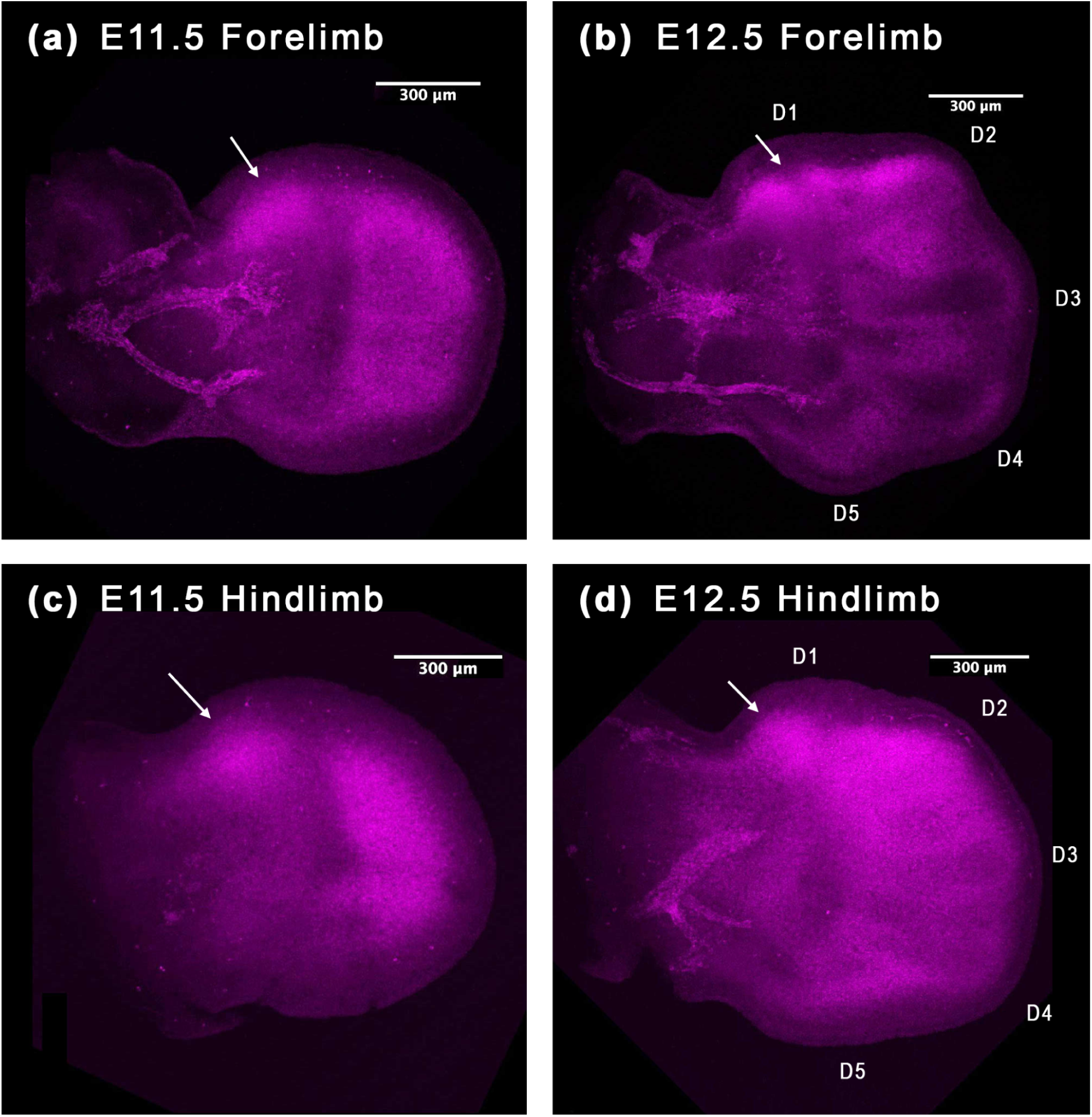
Expression of Lmo4 via hybridization chain reaction (HCR) in mouse E11.5 forelimb (a) and hindlimb (c), and E12.5 forelimb (b) and hindlimb (d). D1-5 = digits 1-5. Anterior = top, distal = right. Scale bar = 300 μm.

### Accelerated evolution of digit 1-loss CREs disrupts binding sites for anteriorly expressed transcription factors

To understand how accelerated evolution of digit 1-loss CREs could impact the mechanisms of limb development, we analyzed the effects of accelerated evolution on transcription factor binding sites for each element. Among the 1,108 thumb-specific elements, binding sites for the transcription factors MEIS1 and MEIS2 were the second and third most frequently lost (Fig. 5a; supplementary table s4). MEIS1 binding sites were lost in 7.67% of the elements and MEIS2 binding sites were lost in 6.86% of elements. MEIS1/2 have been implicated in early anterior-posterior patterning of the limbs, though knockouts of the two genes tend to affect the posterior digits more than the anterior digits (Delgado *et al*. 2021).

**Fig. 5.**
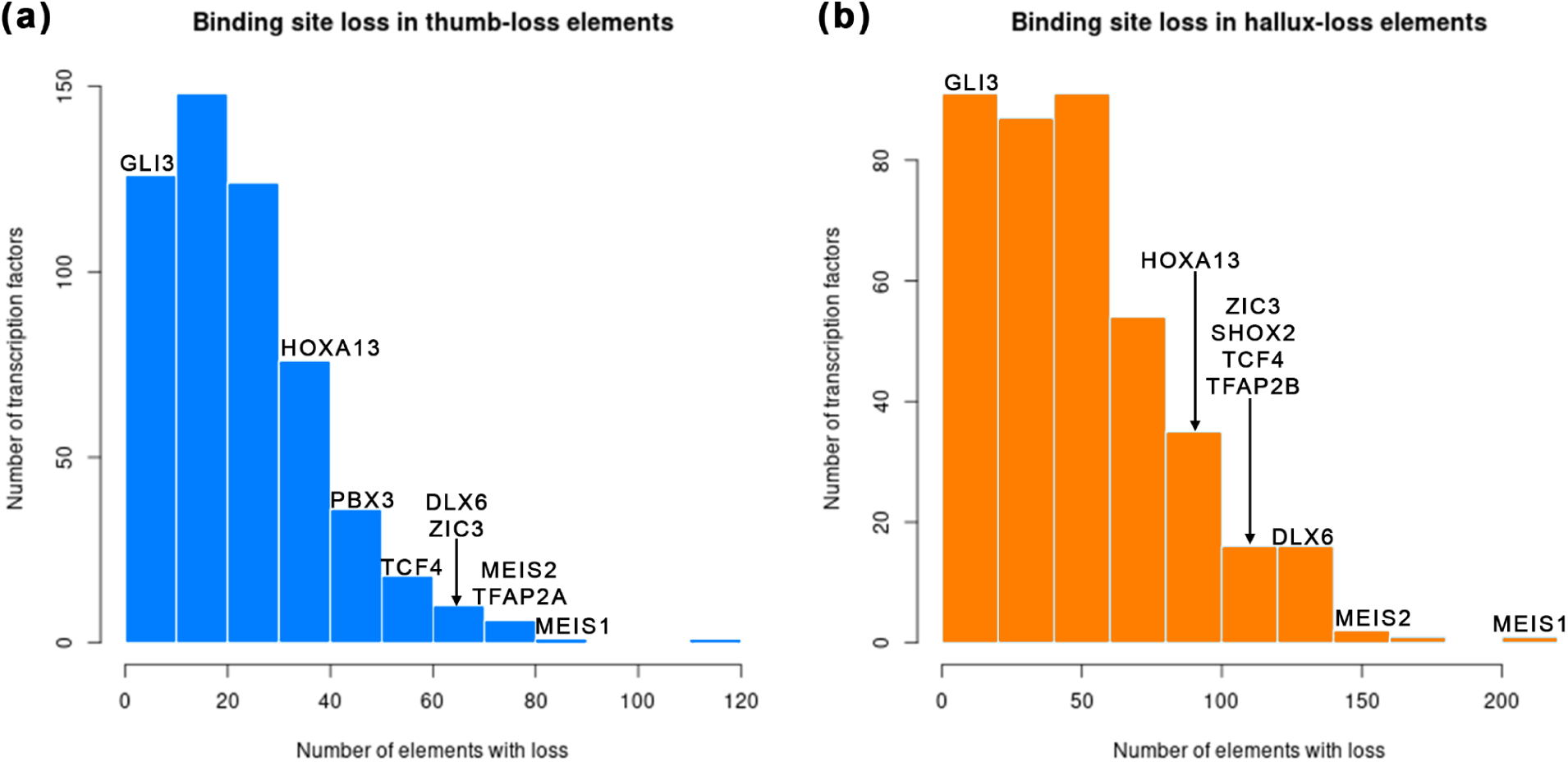
The number of times that binding sites for particular transcription factors have been lost among thumb- (a) and hallux- (b) loss elements. The names of select anterior- and limb-relevant transcription factors are placed above the bins they are contained within.

Overall, anteriorly biased transcription factors are among the most frequently lost binding sites in thumb-loss elements (Fig. 5a, supplementary table s4). DLX6 binding sites were lost in 5.69% of elements, ZIC3 binding sites were lost in 5.42% of elements, and PBX3 binding sites were lost in 3.97% of elements. Binding sites for TFAP2B, which is differentially expressed between digit 1 and the rest of the autopod in a variety of amniotes (Stewart et al. 2019), were lost in 5.14% of elements. *Tcf4* is a forelimb hotspot gene, showing nearby accelerated elements in 6 out of 14 lineages, and we find loss of TCF4 binding sites in 4.78% of elements.

Binding sites for HOXA13 were lost at a high frequency (36 elements; 3.25%). Loss of HOXA13 binding sites in thumb-loss elements contrasts with the lack of accelerated elements near *Hoxa13*. These results suggest that while *Hoxa13* may not be misregulated in the evolution of thumb loss, thumb-relevant transcriptional targets may have lost the ability to interact with HOXA13. GLI3 binding sites were lost in only 3 elements (0.27%), suggesting that GLI3 targets played a minimal role in the evolution of thumb loss.

Analysis of the 2,002 hallux-specific elements uncovered a number of similarities to thumb-loss elements. MEIS1 and MEIS2 binding sites were among the most frequently lost (10.19% and 7.44%, respectively)(Fig. 5b; supplementary table S4). Binding sites for anteriorly biased transcription factors were eliminated from hallux-specific elements at high frequencies, including DLX6 (6.24%), ZIC3 (5.69%), DLX5 (3.55%), and PBX3 (2.75%). As reported above, the regions surrounding each of these anterior genes were determined to be hallux-loss hotspots (**table 1**). Binding sites for proteins encoded by other hallux-loss hotspot genes with known roles in limb development – including *Shox2*, *Tcf4*, *Tfap2b*, *Sox9*, *Hand2*, and *Twist2* – also showed high rates of loss from hallux-specific elements that underwent accelerated evolution. Like thumb-specific elements, hallux-specific elements also show loss of a large number of HOXA13 binding sites (82 elements; 4.10%) but very few GLI3 binding sites (4 elements; 0.20%).

## DISCUSSION

Our results shed light on the genomic trends underlying digit 1 loss. This analysis explains multiple aspects of digit 1’s independence, with implications for both evolutionary and developmental mechanisms.

Our analysis of accelerated non-coding elements suggests overlap between the mechanisms of digit 1 loss in evolution and those involved in digit 1 loss in human congenital disease. This is most readily apparent by the overrepresentation of digit 1-loss elements near genes with known roles in digit 1 loss syndromes. Where evolutionary and disease mechanisms seem to differ is in the types of sequences that are affected around a given gene. For example, our analysis identified cis-regulatory elements near genes involved in human digit loss syndromes, but the latter more commonly originate from exonic perturbation (Van De Laar et al. 2007; Vanlerberghe et al. 2019). If evolutionary loss of digit 1 occurs more frequently via tissue-specific non-coding elements, then this could also explain why species with reduction of digit 1 do not have the range of syndromic phenotypes that result from coding mutations.

Digit 1 loss occurs with two degrees of developmental independence: independence between the forelimb and the hindlimb, and independence from the other four digits. Our results provide some explanation for both of these phenomena.

Regarding independence between the limbs, we found that digit 1 loss elements that are accelerated in a limb-specific manner have stronger associations with human digit loss syndromes, even when the syndromes themselves manifest in both limbs. Thus, even if the associated genes play a role in digit 1 development in both limbs, the expression of these genes may be governed by limb-specific enhancers.

Regarding the independence of digit 1 loss from the other autopodial elements, our data identified accelerated evolution of elements near genes with anteroposteriorly polarized expression patterns in the limb. Some of the most frequent hotspots of acceleration for both thumb loss and hallux loss are around genes known to be expressed in the anterior domains of the limb during the early stages of digit patterning. Furthermore, transcription factor binding sites for many of these same genes are specifically lost at a high rate in digit 1-accelerated elements. Data on the embryonic development of limbs in species with digit 1 loss indicates that the phenotype begins to appear when these genes are expressed.

Our results suggest that digit loss in forelimbs and the hindlimbs involves different mechanisms and genes. Although there is some degree of overlap between forelimb and hindlimb hotspot regions, several anteriorly expressed genes show limb-specific biases in their frequency of evolution. The *Irx3* region seems to be a favored evolutionary locus for thumb loss, showing acceleration in 71% of lineages. No other well-characterized, anteriorly-biased genes were found to be strong thumb-loss hotspots. By contrast, the anteriorly expressed genes *Irx3*, *Sall1*, *Pbx3*, *Dlx5/6*, and *Zic3* are all near hallux-loss hotspots, though none show acceleration in as many lineages as *Irx3* does in thumb-loss lineages. This may suggest that the pathways for evolutionary thumb loss are more constrained than those for hallux loss. Alternatively, hallux development could involve broader reconfiguration of ancestral anterior regulatory networks compared to thumb development.

Despite these limb-specific differences in elements around anteriorly expressed genes, binding sites for anteriorly expressed transcription factors show similar patterns and rates of loss between thumb- and hallux-loss elements. This finding may have implications for the mechanisms by which digit 1 loss evolves. In thumb loss, the broader suite of anteriorly expressed genes need not evolve *cis*-regulatory changes if their target elements have lost the binding sites necessary for interaction. By contrast, hallux loss may have involved disruptions to the regulatory domains of anterior genes and to the binding sites on their targets. As such, it is possible that hallux loss evolved via changes to regulation *of* and *by* anterior transcription factors, while thumb loss was driven mostly by changes to the targets regulated *by* anterior transcription factors.

Prior to this study, some of the most obvious candidates for driving digit 1 loss were *Hoxa13* and *Gli3* (Bastida et al. 2020). Therefore, we were surprised that our analyses did not identify these genes as strong candidates in lineages that evolved thumb-loss or hallux-loss. Neither of these genes were hotspots of accelerated evolution. HOXA13 binding sites were lost in a large number of accelerated elements but did not emerge as an outlier. GLI3 binding sites were almost never lost, implying strong conservation of its role in limb development. This may be because *Gli3* is essential for restricting the effects of *Shh* in multiple anterior components of the limb (Hill et al. 2009). Loss or misregulation of *Gli3* leads to severe polydactyly, perhaps explaining why its regulatory domain and target elements are largely conserved in taxa with digit 1 loss.

Our results also identify potential roles for genes that were not previously considered to be strong actors in digit 1 development. For example, several genes in the Notch signaling pathway – *Hes1*, *Tle3*, *Tle4* – almost always had nearby hallux-specific acceleration. Additionally, *Lmo4* was identified as a strong hotspot that showed acceleration in all but 2 of the thumb-loss and hallux-loss lineages, which led us to investigate its expression and to discover that *Lmo4* is anteriorly biased in the limb. Whether *Lmo4* has a functional role in digit 1 development is yet to be determined, but it is validating to find that our genomic analysis identified genes with relevant expression patterns.

RNA-seq of the developing limbs in digit 1-loss species would significantly expand the findings of this study. Previous research has identified genes involved in bat wing development by comparing RNA-seq of their forelimb with that of their hindlimb. One might imagine a similar approach taken with any species that has lost digit 1 in one set of limbs and not the other, like dogs, cats, or *Jaculus* (jerboa). Such an analysis would not only inform us of which genes are misregulated during digit 1 loss, but could also link misregulation to specific regulatory elements and specific functional mutations within them.

## MATERIAL AND METHODS

### Phenotype identification

Digit 1 phenotypes were identified for 94 species in Zoonomia’s 241 mammal genome alignment (Zoonomia Consortium et al. 2020)(**Table S1**). For most species, digit phenotypes were assigned based on published literature(Van Staden 2014; Senter and Moch 2015; McHorse et al. 2019; Smith et al. 2020) and others were assigned using data from Morphosource (Blackburn et al. 2024) that were corroborated with known phenotypes of other species in the same genus. Species for which the digit phenotype could not be verified or for which there are discrepancies in the literature were omitted from this analysis. For example, cetaceans were omitted from this analysis because there appears to be variability in the number of digit 1 phalanges in many cetacean species (Cooper et al. 2007).

### Identification of digit 1-accelerated CNEs

Multiple alignment format (MAF) files containing these species were extracted from the Zoonomia 241 mammal hierarchical alignment formal (HAL) file (Hickey et al. 2013) using cactus-hal2maf with the following parameters: --filterGapCausingDupes, --noAncestors, --dupeMode single, --refGenome Homo_sapiens, --targetGenomes []. MAF files were extracted separately for each chromosome.

phyloFit (PHAST)(Hubisz et al. 2011) was run on four-fold degenerate sites to generate a nonconserved model of evolution for all species with a known phenotype. phyloFit was run separately on the alignments for each chromosome, and all autosome models were subsequently aggregated using the phyloBoot tool of the PHAST package. The X and Y chromosomes were analyzed separately from the autosomes and from each other throughout this process.

phastCons was run on the alignment for each chromosome to identify genomic regions that are conserved specifically in the species that retain their digit 1 (considered separately for retaining their thumb and retaining their hallux)(parameters: --expected-length 45, --target-coverage 0.3, --rho 0.31, --most-conserved)(Parameter values, excepting --most-conserved, were based on (Nakayama and Makino 2024)). Conserved regions within 5 bp of each other were merged using BedTools (Quinlan and Hall 2010). Elements less than 20 bp or greater than 5000 bp in length were removed from the dataset. Elements that exclusively overlapped exons (as determined by the hg38 annotation) were removed (but elements that only partially overlapped exons were retained).

phyloAcc (Yan *et al*. 2023) was used to identify elements undergoing acceleration in digit 1-loss species. phyloAcc was run separately on thumb- and hallux-conserved elements, with species that have undergone digit reduction used as targets for each analysis. Elements with Bayes factor (BF1) > 10 and Bayes factor (BF2) > 5 were selected as accelerated elements. These accelerated elements were filtered using BedTools to select only elements which overlapped ENCODE’s predicted CREs or ChromHMM predicted limb enhancers during mouse embryonic stages E11.5, E12.5, E13.5, E14.5, or E15.5 (Van Der Velde et al. 2021).

GREAT (McLean et al. 2010) was used to identify genes near digit 1 accelerated elements in hg38 and enriched human phenotypes associated with these elements/genes.

### Identification of thumb/hallux-reduction hotspots

Genomic regions in which more than one third of independent digit 1-loss lineages showed accelerated evolution near a common gene were considered “hotspots” for thumb/hallux reduction. This could include multiple elements around the same gene. Independent thumb-loss lineages are as follows: *Suricata*, *Hyaena*, *Lycaon*, Perissodactyla, *Camelus*, Artiofabula, Caviidae, *Ateles*, *Colobus*, *Tolypeutes*, *Choloepus*, *Trichechus*, Procaviidae, *Orycteropus*. Independent hallux-loss lineages are as follows: *Suricata*, *Hyaena*, Felidae, Canidae, Perissodactyla, Artiodactyla, Lagomorpha, *Thryonomys*, Caviidae + *Dasyprocta*, *Chinchilla*, *Jaculus*, *Dipodomys*, *Choloepus*, *Trichechus*, and Procaviidae.

### Loss of transcription factor binding sites

Transcription factor binding motifs were obtained from the JASPAR CORE Vertebrates non-redundant database (Ovek Baydar et al. 2025) and converted into WTMX format. Data from (Onimaru et al. 2019) was used to determine which transcription factors are expressed in the E11.5 mouse forelimb. Transcription factors with at least one count in each E11.5 replicate and an average of 120 counts at E11.5 were considered to be “expressed” at that stage. For the hindlimb, transcription factors with an average of at least 1 RPKM across all three replicates of E11.5 in (Wang et al. 2018) were considered to be “expressed.” Only transcription factors that are expressed in the E11.5 forelimb or hindlimb (respectively) were analyzed for loss of binding sites. TFforge (Langer and Hiller 2019) was used to identify transcription factor binding sites (TFBSs) that are significantly (p-value < 0.05) lost in digit 1 accelerated elements. The trait-loss list of species was tailored specifically to each element, including only those species which were accelerated for that element.

## REFERENCES

Bastida MF, Pérez-Gómez R, Trofka A, Zhu J, Rada-Iglesias A, Sheth R, Stadler HS, Mackem S, Ros MA. 2020. The formation of the thumb requires direct modulation of *Gli3* transcription by Hoxa13. Proc. Natl. Acad. Sci. 117:1090–1096.

Blackburn DC, Boyer DM, Gray JA, Winchester J, Bates JM, Baumgart SL, Braker E, Coldren D, Conway KW, Rabosky AD, et al. 2024. Increasing the impact of vertebrate scientific collections through 3D imaging: The openVertebrate (oVert) Thematic Collections Network. BioScience 74:169–186.

Carroll SB. 2008. Evo-Devo and an Expanding Evolutionary Synthesis: A Genetic Theory of Morphological Evolution. Cell 134:25–36.

Chiang C, Litingtung Y, Harris MP, Simandl BK, Li Y, Beachy PA, Fallon JF. 2001. Manifestation of the Limb Prepattern: Limb Development in the Absence of Sonic Hedgehog Function. Dev. Biol. 236:421–435.

Cooper LN, Berta A, Dawson SD, Reidenberg JS. 2007. Evolution of hyperphalangy and digit reduction in the cetacean manus. Anat. Rec. 290:654–672.

Delgado I. Control of mouse limb initiation and antero-posterior patterning by Meis transcription factors.

Di Giacomo G, Koss M, Capellini TD, Brendolan A, Pöpperl H, Selleri L. 2006. Spatio-temporal expression of Pbx3 during mouse organogenesis. Gene Expr. Patterns 6:747–757.

Fröbisch NB, Shubin NH. 2011. Salamander limb development: Integrating genes, morphology, and fossils. Dev. Dyn. 240:1087–1099.

Fromental-Ramaint C, Warott X, Messadecq N, LeMeur M, Dolle P, Chambon P. 1995. *Hoxa-13* and *Hoxd-13* play a crucial role in the patterning of the limb autopod. Development 122:2997–3011.

Hickey G, Paten B, Earl D, Zerbino D, Haussler D. 2013. HAL: a hierarchical format for storing and analyzing multiple genome alignments. Bioinformatics 29:1341–1342.

Hubisz MJ, Pollard KS, Siepel A. 2011. PHAST and RPHAST: phylogenetic analysis with space/time models. Brief. Bioinform. 12:41–51.

Jain S, Kim H-G, Lacbawan F, Meliciani I, Wenzel W, Kurth I, Sharma J, Schoeneman M, Ten S, Layman LC, et al. 2011. Unique phenotype in a patient with CHARGE syndrome. Int. J. Pediatr. Endocrinol. 2011:11.

Kawakami Y, Uchiyama Y, Rodriguez Esteban C, Inenaga T, Koyano-Nakagawa N, Kawakami H, Marti M, Kmita M, Monaghan-Nichols P, Nishinakamura R, et al. 2009. Sall genes regulate region-specific morphogenesis in the mouse limb by modulating Hox activities. Development 136:585–594.

Kenny DA, Jurata LW, Saga Y, Gill GN. 1998. Identification and characterization of *LMO 4*, an LMO gene with a novel pattern of expression during embryogenesis. Proc. Natl. Acad. Sci. 95:11257–11262.

Kosicki M, Baltoumas FA, Kelman G, Boverhof J, Ong Y, Cook LE, Dickel DE, Pavlopoulos GA, Pennacchio LA, Visel A. 2025. VISTA Enhancer browser: an updated database of tissue-specific developmental enhancers. Nucleic Acids Res. 53:D324–D330.

Langer BE, Hiller M. 2019. TFforge utilizes large-scale binding site divergence to identify transcriptional regulators involved in phenotypic differences. Nucleic Acids Res. 47:e19–e19.

Li D, Sakuma R, Vakili NA, Mo R, Puviindran V, Deimling S, Zhang X, Hopyan S, Hui C. 2014. Formation of Proximal and Anterior Limb Skeleton Requires Early Function of Irx3 and Irx5 and Is Negatively Regulated by Shh Signaling. Dev. Cell 29:233–240.

Li S-S, Bai S-B, Sun X-F, Yu C-H, Tang Y-N, Jia Z-Q, Li X-P, Shang S-Y, M. Irwin D, Li J, et al. 2025. *Zic3* represses anterior digit development in tetrapods. Zool. Res. 46:684–694.

McHorse BK, Biewener AA, Pierce SE. 2019. The Evolution of a Single Toe in Horses: Causes, Consequences, and the Way Forward. Integr. Comp. Biol. 59:638–655.

McLean CY, Bristor D, Hiller M, Clarke SL, Schaar BT, Lowe CB, Wenger AM, Bejerano G. 2010. GREAT improves functional interpretation of cis-regulatory regions. Nat. Biotechnol. 28:495–501.

Montavon T, Le Garrec J-F, Kerszberg M, Duboule D. 2008. Modeling *Hox* gene regulation in digits: reverse collinearity and the molecular origin of thumbness. Genes Dev. 22:346–359.

Nakayama D, Makino T. 2024. Convergent accelerated evolution of mammal-specific conserved non-coding elements in hibernators. Sci. Rep. 14:11754.

Nguyen JL, Ho CA. 2022. Congenital Disorders of the Pediatric Thumb. JBJS Rev. [Internet] 10. Available from: https://journals.lww.com/10.2106/JBJS.RVW.21.00147

Onimaru K, Tatsumi K, Tanegashima C, Kadota M, Nishimura O, Kuraku S. Developmental hourglass and heterochronic shifts in fin and limb development.:31.

Ovek Baydar D, Rauluseviciute I, Aronsen DR, Blanc-Mathieu R, Bonthuis I, de Beukelaer H, Ferenc K, Jegou A, Kumar V, Lemma RB, et al. 2025. JASPAR 2026: expansion of transcription factor binding profiles and integration of deep learning models. Nucleic Acids Res.:gkaf1209.

Quinlan AR, Hall IM. 2010. BEDTools: a flexible suite of utilities for comparing genomic features. Bioinformatics 26:841–842.

Reno PL, McCollum MA, Cohn MJ, Meindl RS, Hamrick M, Lovejoy CO. 2008. Patterns of correlation and covariation of anthropoid distal forelimb segments correspond to Hoxd expression territories. J. Exp. Zoolog. B Mol. Dev. Evol. 310B:240–258.

Ros MA, Dahn RD, Fernandez-Teran M, Rashka K, Caruccio NC, Hasso SM, Bitgood JJ, Lancman JJ, Fallon JF. 2003. The chick *oligozeugodactyly* (*ozd*) mutant lacks sonic hedgehog function in the limb. Development 130:527–537.

Royle SR, Young JJ. 2021. Developmental biology: A 5ʹHoxd–Gli3 balance in tetrapod axial polarity. Curr. Biol. 31:R1487–R1490.

Senter P, Moch JG. 2015. A critical survey of vestigial structures in the postcranial skeletons of extant mammals. PeerJ 3:e1439.

Sharma D, Mirando AJ, Leinroth A, Long JT, Karner CM, Hilton MJ. 2021. HES1 is a novel downstream modifier of the SHH-GLI3 Axis in the development of preaxial polydactyly.Long F, editor. PLOS Genet. 17:e1009982.

Smith HF, Adrian B, Koshy R, Alwiel R, Grossman A. 2020. Adaptations to cursoriality and digit reduction in the forelimb of the African wild dog (*Lycaon pictus*). PeerJ 8:e9866.

Solounias N, Danowitz M, Stachtiaris E, Khurana A, Araim M, Sayegh M, Natale J. 2018. The evolution and anatomy of the horse manus with an emphasis on digit reduction. R. Soc. Open Sci. 5:171782.

Stearns FW. 2010. One Hundred Years of Pleiotropy: A Retrospective. Genetics 186:767–773.

Stewart TA, Liang C, Cotney JL, Noonan JP, Sanger TJ, Wagner GP. 2019. Evidence against tetrapod-wide digit identities and for a limited frame shift in bird wings. Nat. Commun. 10:3244.

Trofka A, Huang B-L, Zhu J, Heinz WF, Magidson V, Shibata Y, Shi Y-B, Tarchini B, Stadler HS, Kabangu M, et al. 2021. Genetic basis for an evolutionary shift from ancestral preaxial to postaxial limb polarity in non-urodele vertebrates. Curr. Biol. 31:4923–4934.e5.

Van De Laar I, Dooijes D, Hoefsloot L, Simon M, Hoogeboom J, Devriendt K. 2007. Limb anomalies in patients with CHARGE syndrome: An expansion of the phenotype. Am. J. Med. Genet. A. 143A:2712–2715.

Van Der Velde A, Fan K, Tsuji J, Moore JE, Purcaro MJ, Pratt HE, Weng Z. 2021. Annotation of chromatin states in 66 complete mouse epigenomes during development. *Commun*. Biol. 4:239.

Van Staden SL. 2014. THE THORACIC LIMB OF THE SURICATE (*SURICATA SURICATTA*): OSTEOLOGY, RADIOLOGIC ANATOMY, AND FUNCTIONAL MORPHOLOGIC CHANGES. J. Zoo Wildl. Med. 45:476–486.

Vanlerberghe C, Jourdain A-S, Ghoumid J, Frenois F, Mezel A, Vaksmann G, Lenne B, Delobel B, Porchet N, Cormier-Daire V, et al. 2019. Holt-Oram syndrome: clinical and molecular description of 78 patients with TBX5 variants. Eur. J. Hum. Genet. 27:360–368.

Vieux-Rochas M, Bouhali K, Mantero S, Garaffo G, Provero P, Astigiano S, Barbieri O, Caratozzolo MF, Tullo A, Guerrini L, et al. 2013. BMP-Mediated Functional Cooperation between Dlx5;Dlx6 and Msx1;Msx2 during Mammalian Limb Development.Schubert M, editor. PLoS ONE 8:e51700.

Visel A, Minovitsky S, Dubchak I, Pennacchio LA. 2007. VISTA Enhancer Browser--a database of tissue-specific human enhancers. Nucleic Acids Res. 35:D88–D92.

Wang JS, Infante CR, Park S, Menke DB. 2018. PITX1 promotes chondrogenesis and myogenesis in mouse hindlimbs through conserved regulatory targets. Dev. Biol. 434:186–195.

Yan H. PhyloAcc-GT: A Bayesian Method for Inferring Patterns of Substitution Rate Shifts on Targeted Lineages Accounting for Gene Tree Discordance.

Yokoyama S, Furukawa S, Kitada S, Mori M, Saito T, Kawakami K, Belmonte JCI, Kawakami Y, Ito Y, Sato T, et al. 2017. Analysis of transcription factors expressed at the anterior mouse limb bud.Schubert M, editor. PLOS ONE 12:e0175673.

Zhang B, He P, Lawrence JEG, Wang S, Tuck E, Williams BA, Roberts K, Kleshchevnikov V, Mamanova L, Bolt L, et al. 2024. A human embryonic limb cell atlas resolved in space and time. Nature 635:668–678.

Zoonomia Consortium, Genereux DP, Serres A, Armstrong J, Johnson J, Marinescu VD, Murén E, Juan D, Bejerano G, Casewell NR, et al. 2020. A comparative genomics multitool for scientific discovery and conservation. Nature 587:240–245.

